# Ensembled best subset selection using summary statistics for polygenic risk prediction

**DOI:** 10.1101/2023.09.25.559307

**Authors:** Tony Chen, Haoyu Zhang, Rahul Mazumder, Xihong Lin

## Abstract

Polygenic risk scores (PRS) enhance population risk stratification and advance personalized medicine, yet existing methods face a tradeoff between predictive power and computational efficiency. We introduce ALL-Sum, a fast and scalable PRS method that combines an efficient summary statistic-based L_0_L_2_ penalized regression algorithm with an ensembling step that aggregates estimates from different tuning parameters for improved prediction performance. In extensive large-scale simulations across a wide range of polygenicity and genome-wide association studies (GWAS) sample sizes, ALL-Sum consistently outperforms popular alternative methods in terms of prediction accuracy, runtime, and memory usage. We analyze 27 published GWAS summary statistics for 11 complex traits from 9 reputable data sources, including the Global Lipids Genetics Consortium, Breast Cancer Association Consortium, and FinnGen, evaluated using individual-level UKBB data. ALL-Sum achieves the highest accuracy for most traits, particularly for GWAS with large sample sizes. We provide ALL-Sum as a user-friendly command-line software with pre-computed reference data for streamlined user-end analysis.

## Main

Polygenic risk scores (PRS) have emerged as an attractive method for improving risk stratification across a diverse range of traits from cancers to psychiatric disorders^1–6^. PRS can enhance patient risk profiling, refine diagnoses, inform treatment plans, and ultimately improve health outcomes^7–9^. Consequently, PRS has garnered significant interest in both research and clinical practice.

Several methods have been developed to construct PRS based on summary statistics from genome-wide association studies (GWAS). Clumping and thresholding (C+T)-based methods^10,11^, which are fast and widely used, identify a subset of single nucleotide polymorphisms (SNPs) passing thresholds for linkage disequilibrium (LD) and GWAS p-values. However, this filtering process may remove potentially predictive SNPs and ultimately hurt prediction accuracy^12^. Bayesian methods^13–18^ offer intuitive priors to model joints SNP effects and generally strong prediction performance, but their computation is often expensive and sometimes may not even converge, particularly when dealing with large and complex data^18^. Penalized regression-based methods^19,20^ have more efficient optimization algorithms to model joint SNP effects. However, popular penalties such as Lasso can select many false positives under strong correlation^15^, which may lead to reduced prediction accuracy due to complex LD patterns across the genome^21,22^.

In general, these methods require a grid of tuning parameters and identify the best combination for a particular application. While this is a simple way to build a prediction model, it is best suited when one only needs to search through a relatively small tuning parameter space. In high-dimensional problems such as PRS which involve a large number of SNPs, the best tuning parameters may be in the gaps or outside the bounds of the fixed grid being considered^23^. Recent works, in both theory and PRS applications, have demonstrated that ensembling multiple predictors can yield better prediction accuracy than choosing a single best predictor from grid search^24–30^.

In this paper, we propose ALL-Sum <u>A</u>ggregated <u>L</u>0<u>L</u>earn using <u>Sum</u>mary-level data), a fast and scalable method of constructing PRS from publicly-accessible GWAS summary statistics and LD reference. ALL-Sum has two key components. First, we use a L_0_L_2_-penalized regression framework, based on the recently developed L0Learn method^31–33^, that only requires GWAS and LD summary-level data. L0Learn has been shown to better handle complex correlation structures and improve prediction accuracy over methods like Lasso and Bayesian spike-and-slab^32,33^. This offers an attractive penalized approach to model sparse joint effects of SNPs in LD and improve PRS accuracy over methods such as Lassosum2 and LDPred2.

However, a key limitation is that L0Learn requires access to individual-level data, which cannot be easily analyzed in large scale and is often not publicly available. Thus, we modify the optimization problem to only require summary-level data and implement a fast coordinate descent optimization to efficiently provide candidate PRS estimates from a large grid of tuning parameters. Then, in an ensembling step, we use Lasso to estimate weights for each of PRS estimate from different combinations of L_0_ and L_2_ tuning parameters in order to construct a final aggregated PRS. While ensembling typically comes at the practical cost of a dense solution with more SNPs in the final prediction model, it ensures higher prediction accuracy compared to the grid search solution.

We compare ALL-Sum’s performance with four alternative methods – Clumping and Thresholding (C+T)^10,11^, LDPred2^14^, Lassosum2^20^, and Stacked C+T^28^. Variants of these methods have been commonly used in practice^3,34–38^, and we take them to be representative of the broad classes of methods previously discussed. We first evaluate the five methods in large-scale analyses of simulated GWAS data based on European ancestry. Additionally, we construct PRS using 27 published GWAS summary statistics for 11 complex traits from 9 reputable data sources, such as the Breast Cancer Association Consortium^4^, Global Lipids Genetics Consortium^3^, FinnGen^39,40^, and UK Biobank (UKBB)^40,41^. All methods are then validated using independent individual-level data from UKBB. Our simulations and real data analyses show that ALL-Sum can achieve up to 60% improvement in R^2^ for continuous traits and 8% improvement in AUC for binary disease traits, with upwards of 25-times faster runtime and 5-times less memory usage, compared to popular methods.

## Results

### Overview of ALL-Sum framework

ALL-Sum is a computationally efficient PRS method based on summary-level data designed to model joint effects across the genome and improve PRS prediction performance within a specific ancestry (**Methods**). The method consists of two main components: (1) L_0_L_2_ penalized regression using GWAS summary statistics and LD, and (2) aggregation of PRS candidates estimated using different tuning parameter combinations (**Figure 1**). L_0_L_2_ penalized regression allows for control over the number of selected SNPs (L_0_) and shrinkage of effect sizes to avoid overfitting (L_2_). These two penalties together can improve prediction performance by better leveraging SNPs in high LD compared to the L_1_ penalty used in Lassosum, and also have a conceptual connection to the Gaussian spike-and-slab prior commonly used in statistical genetics^13,15,33,42–46^. Using a fast coordinate optimization algorithm, we can efficiently generate effect estimates, with a closed-form at each iteration, given a grid of L_0_ and L_2_ tuning parameter combinations.

**Figure 1.**
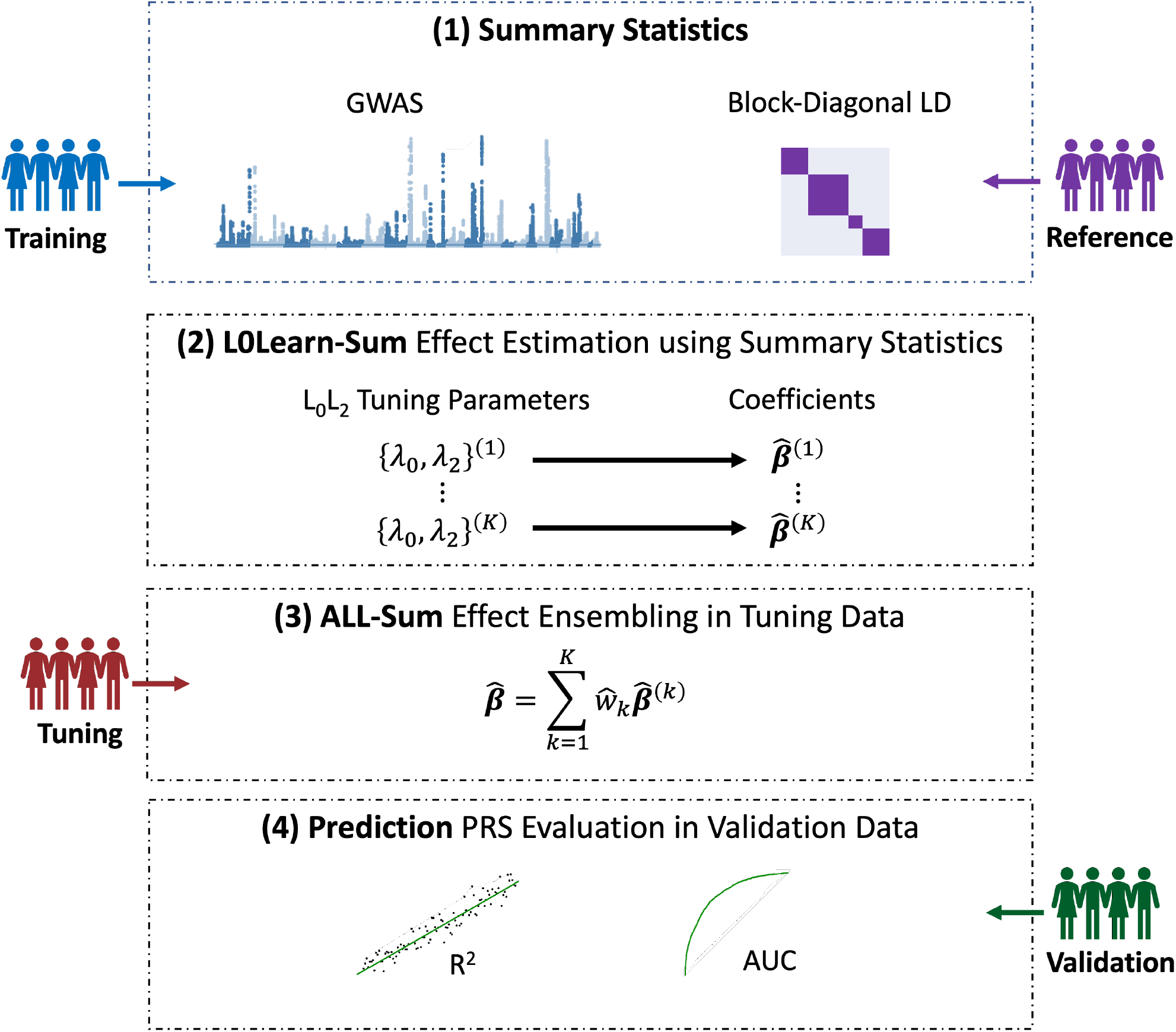
Overview of ALL-Sum framework. (1) Obtain GWAS summary statistics and LD, typically from unobserved training and reference data, respectively. (2) Use the L0Learn-Sum algorithm using GWAS and LD to estimate coefficients 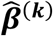 corresponding to L_0_L_2_ tuning parameters {**λ**_1_ **λ**_2_}^(***k***)^ with ***k*** = 1, …, ***K***. (3) Apply each set of tuning parameter-specific coefficients to tuning genotype data to construct ***K*** candidate PRSs. Run cross-validated Lasso regression of the outcome on the ***K*** PRSs to obtain ensembling weights 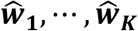 and combine effect estimates from different tuning parameters. (4) Apply ensembled effect estimate to validation data and evaluate final the PRS, using common metrics such as R^2^ or AUC.

These effect estimates are applied to individual-level genotypes in a tuning dataset to construct a set of candidate PRSs corresponding to the tuning parameter grid. Then, in an ensembling step within this tuning dataset, we use the PRSs as predictors in cross-validated Lasso regression to obtain weights for each candidate PRS. The weights are used to aggregate the effect estimates across the tuning parameter grid and to yield a final effect estimate for prediction (**Methods, Figure 1**). In both simulations and real data, we find that an ensemble of multiple candidate PRSs across different tuning parameters leads to more accurate prediction than choosing the single best model from grid search. Furthermore, we show analytically in **Methods** that the classical grid search approach is a special case of the ensembling approach and expected to perform worse than the ensembled estimate.

The default tuning parameters for implementing ALL-Sum include 5 values of λ_2_ between 0.1 and 100 on the log-scale and 50 values of λ_0_ with an adaptive regularization path described in the **Supplementary Method Notes**. Ensembling is done via 3-fold cross-validation lasso to choose 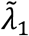 using the defaults of the glmnet package (version 3.0) in R^47^. The implementation of the four alternative PRS methods is provided in the **Methods** section.

### Prediction performance in simulation studies

We first compare prediction performance of ALL-Sum against four alternative PRS methods in a wide range of GWAS sample sizes and causal SNP proportions. Our simulations include 120,000 simulated individuals based on European ancestry generated from HapGen2^48^ and about 1.2 million SNPs from HapMap3^49^. Out of the 120,000 total samples, 100,000 training samples are used to compute GWAS summary statistics, 10,000 tuning samples to compute LD and choose a best PRS model according to each method, and 10,000 validation samples to evaluate out-of-sample prediction accuracy. We vary the causal SNPs from 0.01% to 10% and GWAS sample size from 15,000 to 100,000. For simulation setting, we generate 10 simulation replicates of a continuous phenotype using GCTA^50^ based on a linear additive model with total heritability around 0.4.

Across all sample size and causal SNP proportion settings, we find that ALL-Sum consistently shows top prediction accuracy (**Figure 2, Supplementary Figure 1, Supplementary Table 5**). In the case of extremely low polygenicity (0.01% causal), the signals are strong enough that most methods can get close to the true heritability of 0.4, aligning with previously established bounds on prediction accuracy as a function of causal SNP proportion and sample size^51,52^. Even so, ALL-Sum is able to outperform all methods by about 10% in terms of validation R^2^ on average across all settings (**Supplementary Figure 2**). Under moderately low polygenicity (0.05% and 0.1%) ALL-Sum shows up to 34% in validation R^2^ over LDPred2 and 19% over Lassosum2. As polygenicity increases up to 10% causal, we observe at least around 10% increase in R^2^ by ALL-Sum over Stacked C+T and Lassosum2. Across all settings, ALL-Sum vastly outperforms simple C+T, achieving up to twice the R^2^ in some settings. Alternative methods seem to show preference to either very strong (Stacked C+T and Lasossum2) or weak (LDPred2) signals with respect to both polygenicity and GWAS sample size.

**Figure 2.**
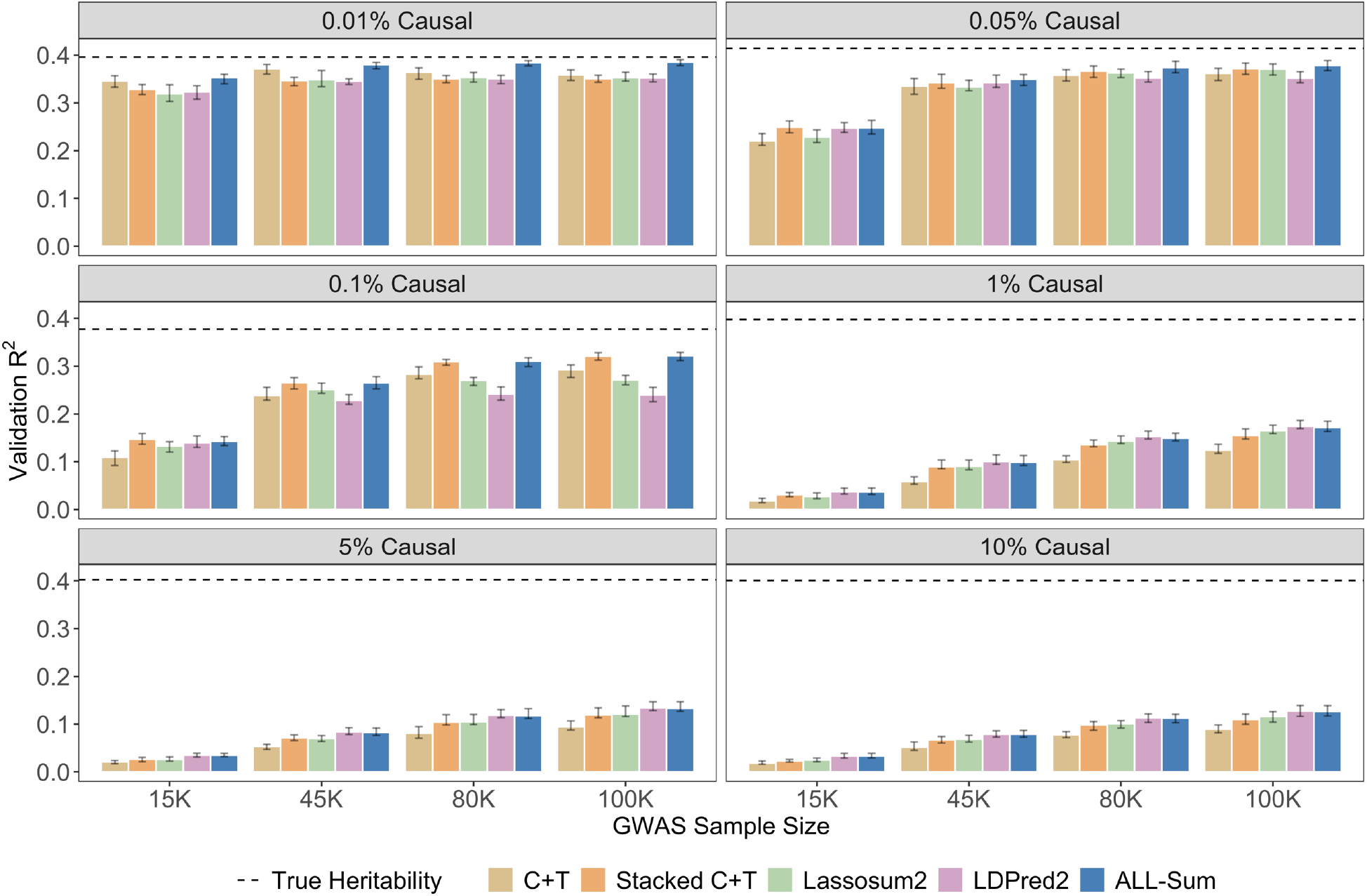
Method comparison in simulation studies with 1.2 million SNPs with varying proportions of causal SNPs. The predictive accuracy of ALL-Sum and four alternative PRS methods are evaluated on simulated data across six causal SNP proportions (panels) and four GWAS sample sizes (X-axis). The Y-axis corresponds to R^2^ on 10,000 validation samples set after choosing a best PRS for each method on 10,000 tuning samples. Bar plots indicate the average R^2^ across 10 simulation replicates for each setting, and error bars represent the range in R^2^.

However, leveraging both the L_0_L_2_ regression and ensembling steps allows ALL-Sum to perform well in all scenarios.

### Runtime and memory usage in simulation studies

Beyond prediction performance, we also evaluate the runtime and memory usage of ALL-Sum on three chromosomes (**Figure 3, Supplementary Table 6**). We use LDPred2 as a benchmark for comparison, given its widespread usage in the field^34,53^.

**Figure 3.**
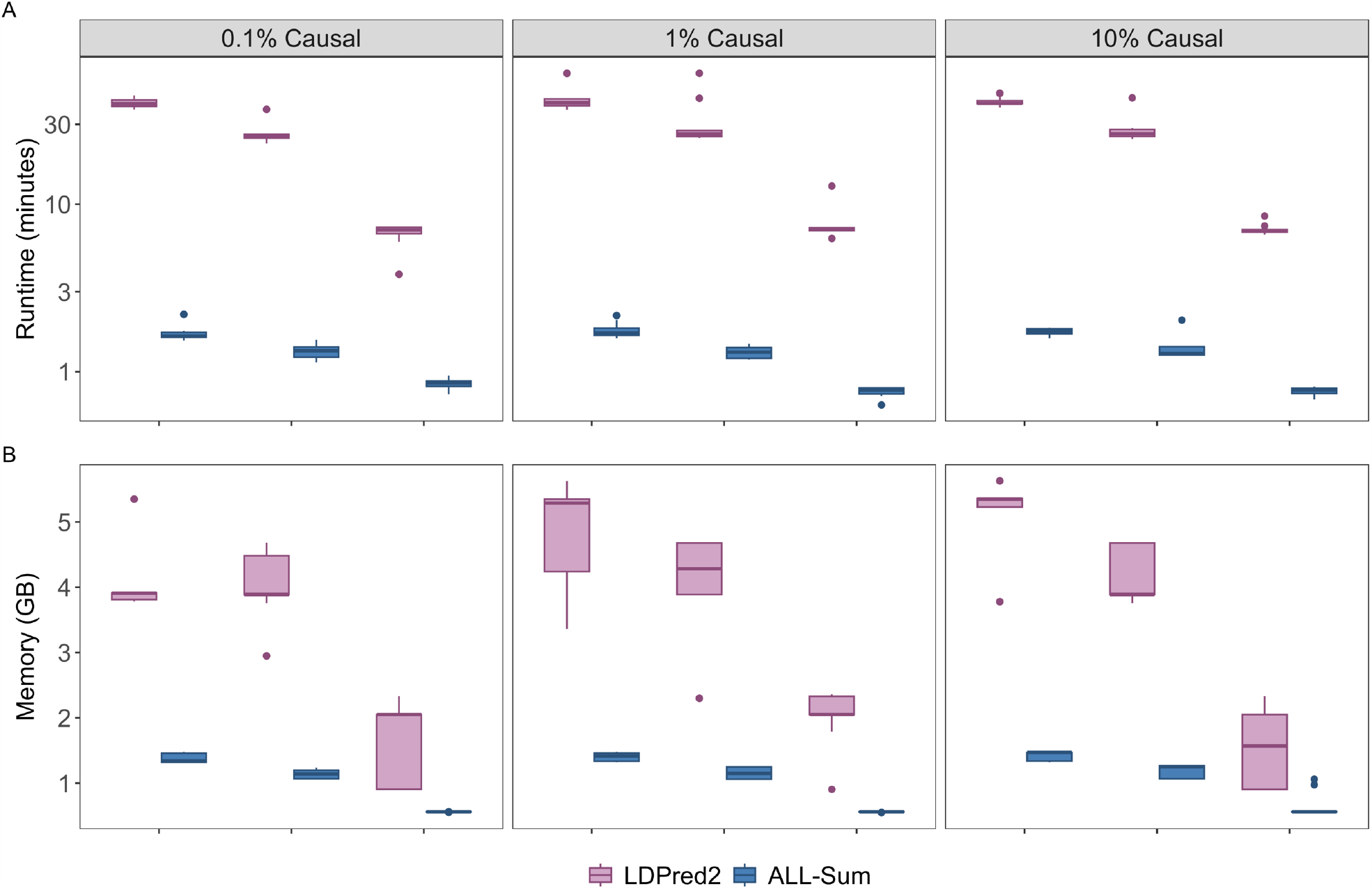
Runtime and memory usage comparison between ALL-Sum and LDPred2. Runtime (Y-axis A) and memory usage (Y-axis B) were compared using simulated data on chromosomes 1, 10, and 21 (X-axis), with causal SNP proportions 0.1%, 1%, and 10% (panels) and GWAS sample size of 100,000.

Across all settings, ALL-Sum runs about 20 times faster than LDPred2 on average for larger chromosomes while using up to four times less memory (**Supplementary Figure 3**). This efficiency can be attributed to the rapid coordinate descent algorithm, and the integration with plink 2.0 for efficiently computing multiple risk scores at once^54^. Within the ALL-Sum workflow, most of the runtime for ALL-Sum is dedicated to coordinate descent, while a consistent proportion of time is taken up by computing PRS at various steps (**Supplementary Figure 4**). It is important to note that LDPred2 and other methods implemented in the bigsnpr R package (Stacked C+T and Lassosum2), not only require memory during analysis but also when generating new backing files that require substantial additional memory on top of the existing plink-format genotype data^55^.

### Continuous trait prediction in UK Biobank

To evaluate the practical usage of ALL-Sum, we apply ALL-Sum and the four alternative methods using large-scale consortium- and biobank-based GWAS summary statistics (**Supplementary Tables 1-4**). We perform tuning and validation of the PRS methods using genotype and phenotype data of 40,000 randomly selected European samples from the UK Biobank (UKBB), with 20,000 samples used for each step. We used publicly available GWAS for 6 continuous traits: high-density lipoprotein (HDL) cholesterol, low-density lipoprotein (LDL) cholesterol, log-triglycerides (logTG), and total cholesterol (TC) from the Global Lipids Genetics Consortium (GLGC), as well as height and body mass index (BMI) from the Genetic Investigation of ANthropometric Traits (GIANT) Consortium. We specifically chose summary statistics that exclude UKBB samples to ensure valid prediction evaluation. To provide a broader comparison, we use the remaining 278,052 European UKBB samples not used in tuning or validation as additional training data to compute GWAS summary statistics for each trait.

Across nearly all traits and data sources, ALL-Sum shows the best prediction performance, measured by R^2^ of the PRS in the 20,000 validation samples adjusted for age, sex, and the first 10 principal components (PCs) (**Figure 4, Supplementary Table 7**). ALL-Sum achieves up to 60% higher R^2^ compared to alternative methods (**Supplementary Figure 5**). We see particularly strong performance of ALL-Sum using the GLGC-based summary statistics, which have about four times larger sample size than UKBB, suggesting that our method can make better use of well-powered GWAS than other methods. The GIANT- and UKBB-based summary statistics have similar sample sizes, but the GIANT summary statistics contain fewer SNPs after filtering to overlap with the UKBB genotype data. In this case, we notice that ALL-Sum performs better using the UKBB summary statistics, showing the increased predictive potential with a larger set of SNPs to analyze.

Within LDPred2 results, we find that prediction accuracy tends to be better using UKBB-based GWAS summary statistics regardless of differences in sample size or number of SNPs, suggesting a sensitivity to the similarity in underlying genetic architecture between the data used to compute GWAS summary statistics and LD, which has also been observed in previous analysis^20^. Furthermore, LDPred2 failed to converge after trying 20 different random seeds for HDL using GLGC summary statistics, demonstrating practical implementation issues with the MCMC algorithm.

**Figure 4.**
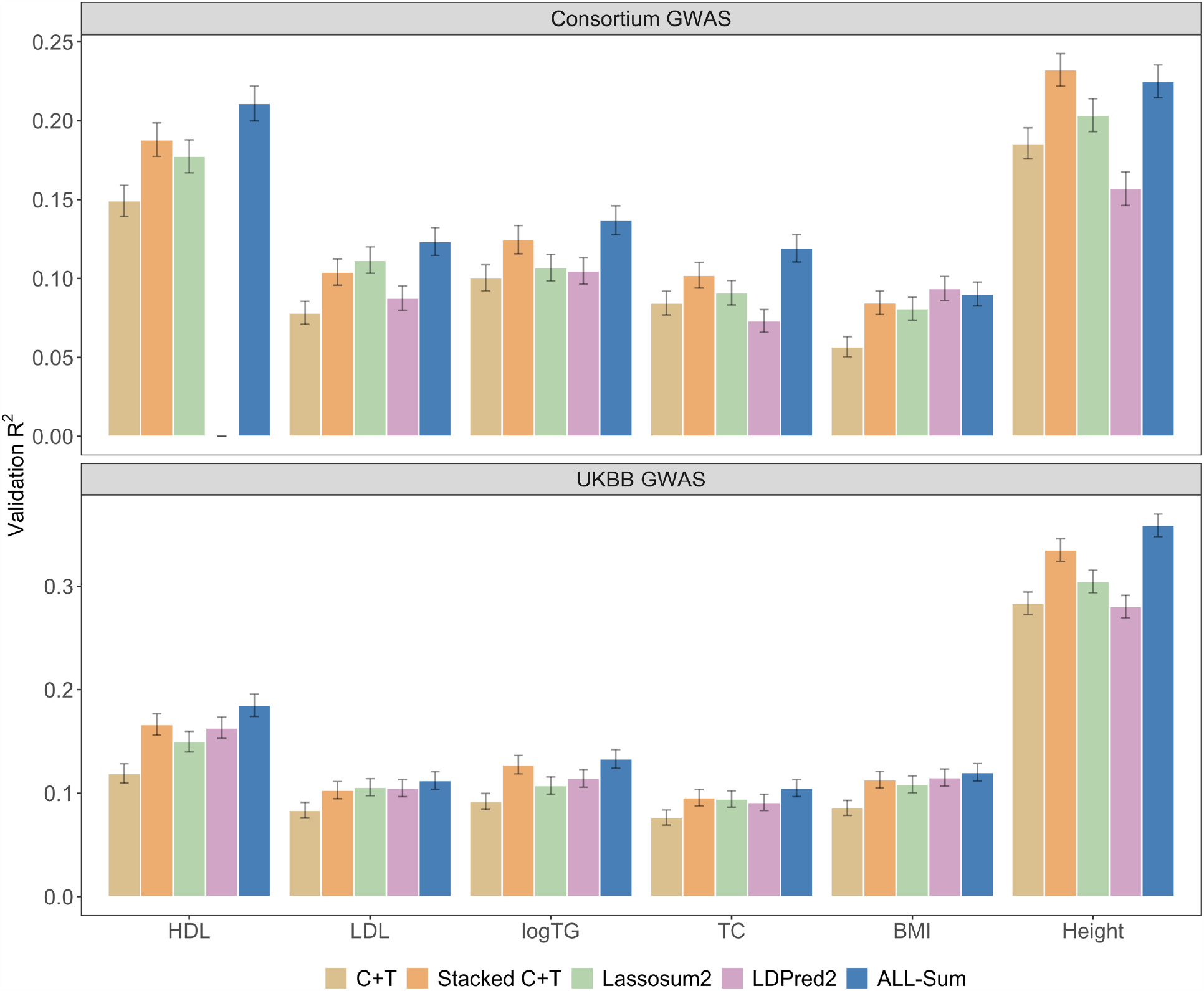
Prediction accuracy comparison of PRS for continuous traits using consortium- and UKBB-based GWAS, validated in UKBB. We compare the prediction accuracy (Y-axis) of ALL-Sum and 4 alternative PRS methods on 6 continuous traits (X-axis), train using summary statistics from either consortium-based (top panel) or UKBB-based GWAS (bottom panel) with sample sizes up to 930,000 and 276,000, respectively. Prediction accuracy is measured as the R^2^ of the PRS from each method with respect to the residuals of a trait after regression out age, sex, and the first 10 PCs. Bar plots indicate the bootstrap mean R^2^ over 10,000 resamples within a validation sample of 20,000 European individuals in the UKBB, independent from those used for UKBB-based GWAS. Error bars indicate the bootstrap 95% confidence interval. (Note: LDPred2 failed to converge for HDL using consortium GWAS summary statistics)

### Disease risk prediction in UK Biobank

We further apply the PRS methods to five binary disease traits: asthma, coronary artery disease (CAD), type 2 diabetes (T2D), breast cancer, and prostate cancer. We obtain summary statistics from case-control GWAS published by reputable consortia: Transnational-National Asthma Genetic Consortium (TNAGC)^56^, Coronary Artery Disease Genome-wide Replication and Meta-analysis plus The Coronary Artery Disease Genetics (CARDIoGRAMplusC4D)^57^, Diabetes Genetics Replication And Meta-analysis (DIAGRAM) consortium^58^, Breast Cancer Association Consortium (BCAC)^4^, and Prostate Cancer Association Group to Investigate Cancer Associated Alterations in the Genome (PRACTICAL)^59^. Additionally, we obtained GWAS summary statistics from FinnGen^39^, which published several well-powered binary disease GWAS, and the 278,052 European UKBB samples. Using the same analysis workflow as with continuous traits, we evaluate each PRS method’s performance using Area under the ROC Curve (AUC) adjusted for baseline covariates^60^.

Across various disease traits and data sources, ALL-Sum again shows generally better and robust performance. Compared to alternative methods, ALL-Sum shows up to 8% increase in AUC over alternative PRS methods (**Supplementary Figure 5**) and typically achieves either the highest or second-highest AUC across all methods (**Figure 5, Supplementary Table 8**) and. ALL-Sum particularly excels in predicting T2D and prostate cancer, whereas LDPred2 performs well for CAD and Breast Cancer. As expected from continuous trait analyses, ALL-Sum generally shows significant prediction gains when using consortium-based GWAS that have a large number of both cases and controls. Meanwhile, UKBB generally has few cases, resulting in potentially underpowered summary statistics. While FinnGen has fairly well-powered summary statistics relative to UKBB, but genetic differences between the Finnish and British populations may explain why there is not a substantial improvement in prediction accuracy^39,61^.

**Figure 5.**
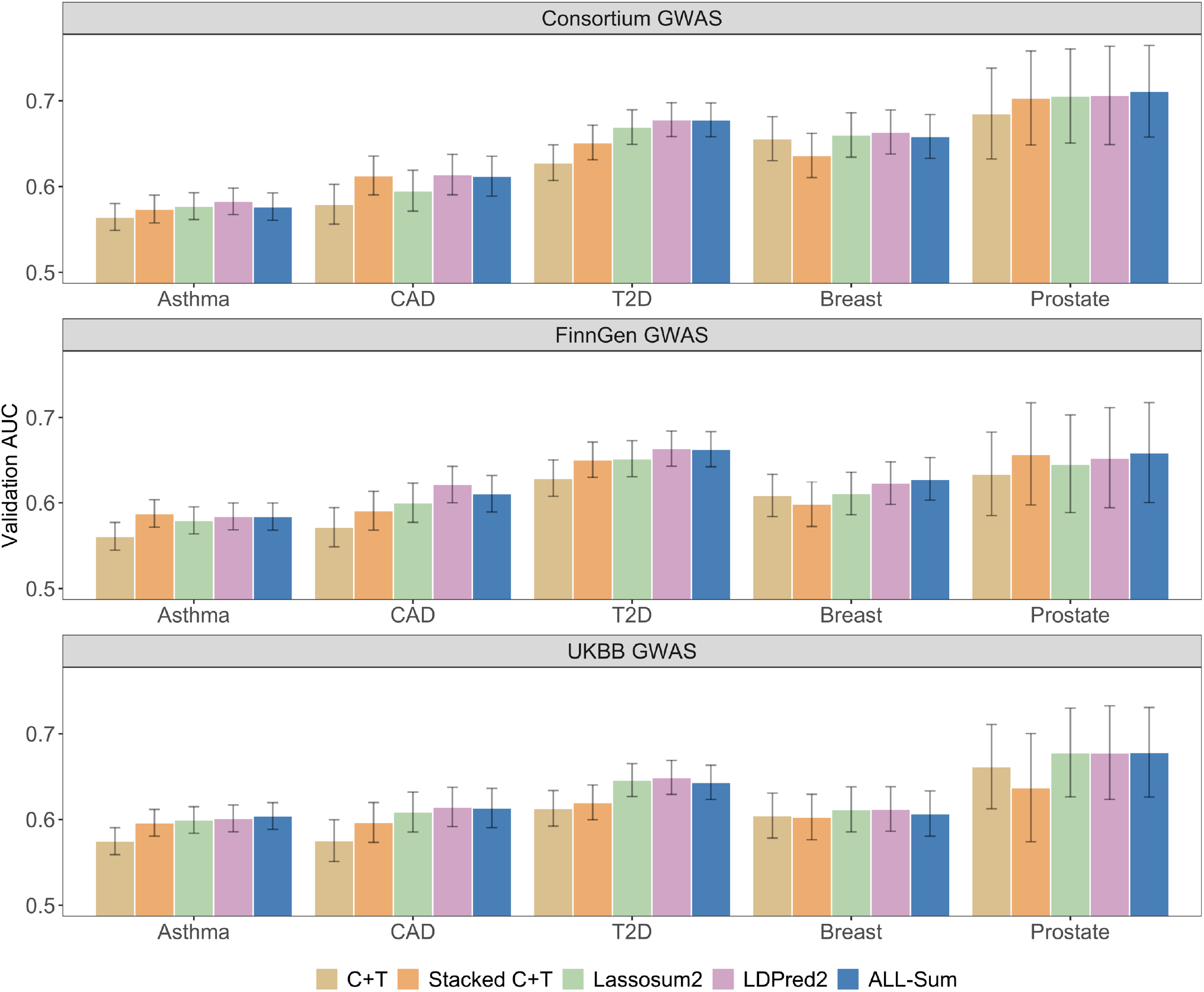
Prediction accuracy comparison of PRS for binary traits using consortium-, FinnGen-, and UKBB-based GWAS, validated in UKBB. We compare the prediction accuracy (Y-axis) of ALL-Sum and 4 alternative PRS methods on 6 continuous traits (X-axis), using summary statistics from either consortium-based (top panel), FinnGen-based (middle panel), or UKBB-based GWAS (bottom panel) with up to 180,000, 49,000, and 20,000 cases, respectively. Prediction accuracy is measured as the adjusted AUC of the PRS from each method conditional on age, sex, and the first 10 PCs. Sex is not used as a covariate for breast and prostate cancers. Bar plots indicate the bootstrap mean R^2^ over 10,000 resamples within a validation sample of 20,000 European individuals in the UKBB, independent from those used for UKBB-based GWAS. Error bars indicate the bootstrap 95% confidence interval.

### Runtime and memory usage for prediction in UK Biobank

To further verify the practical usability of our method, we also compare the runtime and memory usage of ALL-Sum and LDPred2 using the 11 full UKBB-based summary statistics (**Supplementary Figure 6, Supplementary Table 9**). We find that ALL-Sum completes full analysis, from processing input data to final validation, in less than 30 minutes and using about 20 GB for both continuous and binary traits. On the other hand, LDPred2 takes at least a few hours to run without parallelization, with up to 25 times longer runtime and 3 times more memory compared to ALL-Sum. In these real data settings, ALL-Sum consistently requires far fewer computational resources to deliver similar to better risk stratification than popular Bayesian computing.

### Prediction gains from ensembling over grid search of tuning parameters

As a key component of our method is the ensembling step that aggregates PRS models estimated using different tuning parameters to improve prediction accuracy and stability over a simple grid search for the optimal tuning parameters. Our simulation studies reveal that in the extreme high and low polygenicity settings, grid search and ensembling yield similar predication accuracy (**Supplementary Figure 7, Supplementary Table 10**). With low polygenicity, both estimates can identify a highly predictive PRS, given the strength of the signals^51,52^. At high polygenicity, weak and dense signals are adequately captured by the grid search estimate since the L_2_ penalty component yields a dense ridge-like estimate. However, at the intermediate polygenicity levels, ensembling offers a more noticeable gain in R^2^, suggesting that the tuning parameter grid may not always contain the best parameter for the grid search PRS estimate to perform well. Ensembling ensures high prediction accuracy by borrowing information from multiple PRS candidates, but it also includes a larger number of SNPs as a trade-off. For practical applications that require a PRS with few SNPs, grid search may be more suitable, despite the additional fine-tuning required.

To further evaluate our ensembling approach, we compare the prediction performance of ensembling with Lasso (L_1_-penalty) against standard grids search, as well as ensembling with Ridge (L_2_-penalty). Our experiments on real continuous traits favor Lasso for ensembling different PRS estimates (**Supplementary Figure 8, Supplementary Table 11**). Ensembling with Ridge can achieve similar prediction accuracy as Lasso across all traits. However, since no variable selection is performed, Ridge keeps all candidate PRS and leads to a much larger number of selected SNPs in the final prediction model. Thus, while ensembling will yield a much denser solution than grid search, we choose to use Lasso since it still performs some selection to prioritize the most predictive candidate PRS.

### Stability across different LD references

We further evaluate ALL-Sum’s performance using different LD references, given that GWAS summary statistics and LD are typically computed on different genotype data. We assess the validation R^2^ in both simulations and UKBB continuous traits with three LD reference panels: the training data used for GWAS, the tuning data used in our primary analyses, and the 503 European samples from the 1000 Genomes Project (1000G)^62^.

Our analyses using both simulated and UKBB-based lipid summary statistics show virtually identical results across when using either the training or tuning data as the LD reference (**Supplementary Figure 9-10, Supplementary Table 12-13**). There is a slight drop in prediction accuracy when using UKBB-based summary statistics and 1000G-based LD, about 4% lower R^2^ on average compared to using LD based on the UKBB tuning data. Although both UKBB and 1000G LD only consist of European ancestry, the UKBB is primarily British Europeans, while 1000G includes a wider range of European populations.

Our analyses suggest that ALL-Sum is fairly robust to the use of different LD references. Specifically, prediction accuracy shows negligible differences when using in-sample LD from training data or external LD from tuning or 1000 Genomes data. While prediction accuracy is better when using UKBB-based LD for UKBB-based GWAS, using 1000G-based LD only diminishes prediction accuracy by a small percentage and is still about the same if not better than other methods using UKBB-based LD in the main analysis. Furthermore, we show that using LD based on tuning data yields the same prediction as using LD based on the training data used for generating GWAS summary statistics. Thus, we can just use LD computed from a small and accessible dataset, such as the tuning data or 1000G, so long as the population is similar to that of the GWAS. As part of our software package, we provide two pre-computed LD based on 1000 Genomes Project and UKBB for convenient direct input for analysis.

## Discussion

In this paper, we propose ALL-Sum as a fast and scalable PRS method that combines efficient L_0_L_2_ optimization using publicly accessible GWAS summary statistics and LD with ensembling of PRS estimates from different tuning parameters to improve prediction accuracy. Extensive simulation studies and real data analyses show that our method consistently provides strong prediction performance across a broad spectrum of traits and data sources, is robust to different LD reference, and uses much less time and memory compared to the state-of-the-art Bayesian approach. ALL-Sum’s performance in improving PRS prediction is most evident in continuous traits and GWAS with large sample sizes. The prediction performance for binary traits is more variable due to the predominance of healthy individuals in UKBB and hence few disease cases^63^. However, with the ongoing expansion of biobank data, we anticipate the potential for richer clinical data and more accurate disease risk prediction in the future.

Although ALL-Sum introduces improvements in prediction and computational and resource efficiency, there are several limitations and opportunities for continued work. The current model is designed for single ancestry analysis. While a large portion of GWAS data currently comes from individuals of European ancestry, it is well-known that polygenic risk scores do not transfer well between individuals of different ancestries, which can impact their utility for patients of non-European ancestry^64–69^. Many methods that utilize summary data from multiple populations have already been proposed and demonstrate improved prediction in under-represented populations^25,29,30,70–74^. As genetics research moves towards greater diversity, ALL-Sum serves as a valuable foundation for extension to incorporating data from multiple ancestries, as well as admixed individuals^75–77^.

Our coordinate descent algorithm significantly reduces runtime compared to Bayesian computation, but there are several directions primed for further software refinement. One way is to devise a smaller data-driven tuning parameter grid similar to the “auto” functions in some Bayesian softwares^14,16^, which would reduce calls to coordinate descent algorithm and reduce both runtime and memory usage. Future work can also incorporate active set updates and combinatorial swaps from the original L0Learn algorithm, as well as parallelization and sparse matrices, to speed up optimization and potentially give better prediction accuracy^31,78,79^. In terms of ensembling, we focus on Lasso given its well-established software and defaults in the glmnet R package, simple linear interpretations, and implementation in other ensembling contexts^28^. Future work can incorporate more intricate strategies, such as those provided in the SuperLearner package^80^, to provide even more flexible models to further improve prediction accuracy.

Finally, while the first step of L_0_L_2_ penalized regression is applied to summary-level data alone to construct PRS models, we still need access to individual-level data for ensembling within the tuning dataset. This is also the case for tuning and validation of the other methods compared in this paper. Having sufficient data for both tuning and validation may be difficult to come by, especially for rare binary outcomes that are not well-recorded in well-established sources like UKBB^63^. Furthermore, the cross-validation step for ensembling may be sensitive if there are few samples or cases in the tuning dataset, although this will similarly affect other methods. Methods have proposed for tuning PRS methods without individual-level data^81–85^. However, final prediction accuracy in final validation may not be as good due to approximations either from generating proxy new data or using unsupervised approaches to the best PRS.

To conclude, we introduce ALL-Sum as a fast and scalable method for PRS using summary-level data and demonstrate its strong prediction performance through comprehensive evaluations in simulations and real data analysis. ALL-Sum is available as a streamlined R software from automated data processing to final validation, using simple command-line inputs.

## Acknowledgements

We would like to thank Jin Jin, Jingwen Zhang, Hussein Hazimeh, Hui Li, Peter Kraft, and Rui Duan for their assistance, insights, and feedback on this project. We are grateful to participants in the UK Biobank (obtained under UK Biobank resource application 52008), FinnGen, and all consortiums referenced in this paper for providing data necessary for our analysis. All analyses were conducted on Harvard FAS Research Computing. This research was supported by NIH Training Grant T32GM135117 and NSF Graduate Research Fellowship DGE-2140743 (T.C.), NIH grants K99 CA256513 and NIH Intramural Research Program (H.Z.), Office of Naval Research grants N000142112841 and N000142212665 (R.M.), and NIH grants R35-3 CA197449, U19-CA203654, R01-HL163560, U01-HG009088 U01-HG012064 (X.L.).

## Author Contributions

All four authors conceived the project. T.C. carried out the data analyses and developed the software and online resources for sharing, under the supervision of H.Z., R.M., and X.L. All authors drafted, reviewed, and approved the final version of this manuscript.

## Data and Code Availability

Simulated genotype data can be found on the Harvard Dataverse (https://dataverse.harvard.edu/dataverse/multiancestry). Access to FinnGen- and consortium-based GWAS summary statistics are detailed in **Supplementary Table 1**. The ALL-Sum software is available at https://github.com/chen-tony/ALL-Sum/. The repository includes the main R and C++ code for analysis, as well as a tutorial with example data. Pre-computed European-based LD from the 1000 Genomes Project and UKBB will be available on the Harvard Dataverse (https://dataverse.harvard.edu/dataverse/allsum).

## Methods

### ALL-Sum

#### L0Learn-based prediction using individual-level data

Suppose we have a linear model

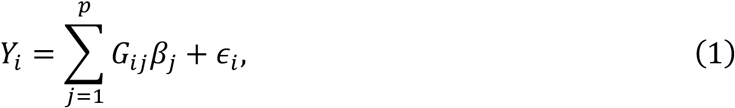

where *Y*_*i*_ is the observed phenotype for an individual *i* ∈ {1, … *n*}, *β*_*j*_is the effect size for SNP *j* ∈ {1, …, *p*}, *G*_*ij*_is the dosage of SNP *j* for individual *i* that takes values between 0 and 2, and *ϵ*_*i*_ is a normally distributed error term.

In a high-dimensional setting where *p* can be several orders of magnitude larger than *n*, we assume that a subset of SNPs are causal, or with nonzero effects *β*_*j*_. This motivates the use of penalties to encourage sparsity in our effect estimate^86–88^. L0Learn was recently proposed as a general penalized regression framework that directly penalizes the number of nonzero effects to give very sparse estimates^31^. Specifically, the method aims to estimate the regression coefficients ***β*** = (*β*_1_, …, *β*_*p*_) by minimizing the penalized least squares loss function:

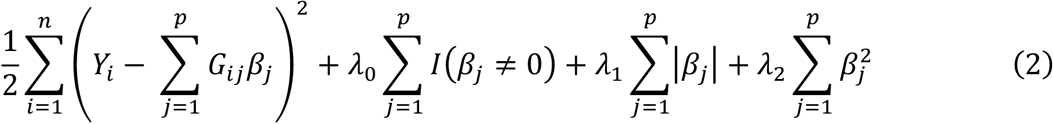

The first term is the squared error from using ***β*** as coefficients for each SNP. In the second term, *I*(*β*_*j*_ ≠ 0) is an indicator that equals 1 if the *j*-th entry of ***β*** is nonzero and 0 otherwise. The sum of these indicators forms the L_0_ pseudonorm, which describes the sparsity of ***β*** by the total number of nonzero entries. The parameter *λ*_0_ controls the strength of the penalty on nonzero entries, where higher values impose stronger penalties and thus fewer nonzero entries. The third and fourth terms refer to additional regularization on either the absolute or squared value of *β*_*j*_. The corresponding penalty parameters *λ*_1_ and *λ*_2_ enforce shrinkage on the estimated effect sizes, reducing overfitting to the training data and improve out-of-sample prediction.

#### L0Learn-based prediction using GWAS summary statistics

The loss function in (2), which was used in the original L0Learn method^31^, requires individual-level data that often cannot be easily shared publicly. Furthermore, the algorithm may not scale easily to the order of genome-scale analysis, involving millions of SNPs and hundreds of thousands of individuals. To address these issues, we first propose L0Learn using publicly accessible LD and GWAS summary-level statistics. Specifically, equation (2) can be rewritten more concisely into matrix notation as:

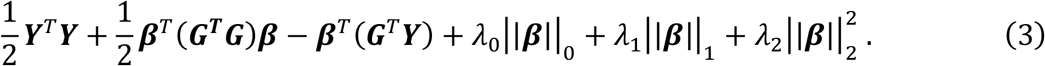

Assuming ***Y*** and each column of ***G*** are standardized to have mean 0 and variance 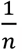, we can transform equation (3) to the following equation, which only depends on GWAS association summary statistics and the LD matrix:

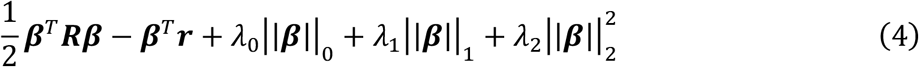

***R*** is a *p × p* matrix representing the LD matrix approximating ***G***^*T*^***G*** when the genotypes are standardized. ***r*** is a *p*-length vector corresponding to ***G***^*T*^***Y***, which is equal to the correlation of each SNP with the phenotype under the standardization. Each entry *r*_*j*_can be computed as 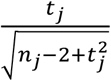, where 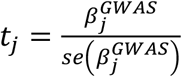 is the GWAS test statistic for the marginal SNP effect and *n*_*j*_is the sample size observed for that SNP. This formula for *r*_*j*_is based on the test statistic for Pearson’s correlation and is nearly identical to 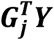 for large *n*_*j*_.

#### Scalable coordinate descent optimization

The optimization problem can be solved using cyclical coordinate descent, an efficient algorithm for high-dimensional problems^31,47^. Given an initialization ***β***^(0)^, each coordinate or variable *j* at timestep *t* is iteratively updated by the following rule, which has a closed form:

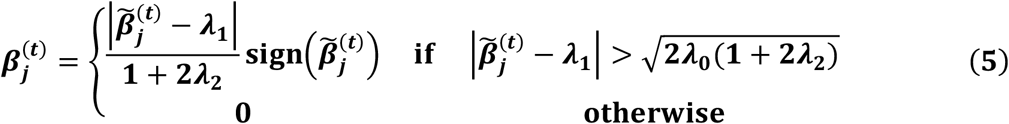

where 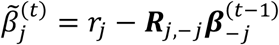 represents the residualized marginal effect of SNP *j*, adjusting for effects of LD by neighboring SNPs. ***R***_*j*,-*j*_refers to the *j*-th row of *R* after removing the *j*-th column entry. Similarly, 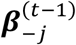 removes the *j*-th entry of ***β***^(*t*-1)^. In the context of sharing and accessing large-scale genetic data, ***r*** and ***R*** are typically computed from different sets of genotypes. For instance, ***r*** may come from large publicly-available GWAS, and ***R*** from well-established reference data such as the 1000 Genomes Project^62^. For the L_0_L_2_-penalized estimator, equation (5) simplifies to

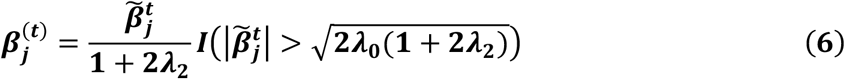

where *I*(⋅) is an indicator function that equals 1 if the inequality in parentheses is satisfied and 0 otherwise.

In practice, using the entire LD matrix is far too large to store and share. Furthermore, since the number of SNPs often far exceeds the number of samples, estimating all pairwise correlations can be noisy and potentially hinder downstream analysis^89,90^. A simple workaround is to approximate LD by a sparse matrix with only a subset of pairwise correlations. Our current implementation focuses on a block-diagonal approximation using LD blocks based on published base-pair positions corresponding to roughly independent regions along the genome^41^. This simplification greatly improves the efficiency of the algorithm by effectively dividing the overall optimization problem into smaller independent problems.

Since the penalized regression problem assumes that both the outcome and genotypes have all been standardized, we can re-scale the effect estimates so that they correspond to genotypes on the original scale, where SNP dosage ranges from 0 to 2. This adjustment requires access to minor allele frequencies (MAF), which may either be included with the GWAS summary statistics or estimated using available genotype data. For a given effect estimate 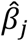and corresponding SNP minor allele frequency *π*_*j*_, the rescaled effect estimate is given as 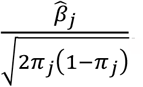.

#### Ensembled PRS from tuning parameter grid

After producing multiple estimates for ***β*** according to a grid of tuning parameter combinations, the immediate next step is to choose the ***β*** that gives the best fit on a tuning dataset. Traditional grid search for fine-tuning is simple and may work well if the best tuning parameters can be easily identified within the grid. However, this may miss out on the parameterization that yields the best predictor. First, the optimal tuning parameters may lie outside the fixed grid. Second, more importantly, as shown in equations (4) and (5), the same tuning parameters are used for all SNPs. For example, in equation (6), at each iteration the same hard threshold 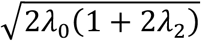 and shrinkage 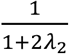 are used for estimating all coefficients *β*_*j*_.

To identify a high-accuracy and more flexible prediction model, we propose an ensemble method that combines different 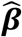 across the whole tuning parameter grid. This is done by estimating an optimal weighting using cross-validated Lasso regression. Ensembling across the tuning parameter grid allows different *β*_*j*_to follow different thresholding and shrinkage.

Specifically, suppose we have a grid of *N* different combinations of tuning parameters *λ*_0_ and *λ*_2_. For each grid point *k* ∈ {1, … . *K*}, we generate effect estimates 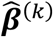 to tuning parameters 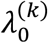 and 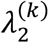. We first multiply each effect estimate by the genotypes in a tuning dataset to compute PRS 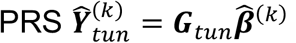. These PRS are then used as covariates in a Lasso regression of the outcome in the tuning dataset ***Y***_*tun*_ on the PRSs 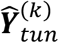 to estimate the weights *w*_*k*_ by minimizing the penalized regression

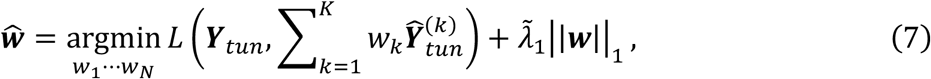

where 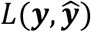 is either the least squares loss 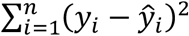 for continuous outcomes or logistic loss 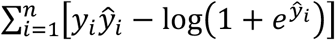 for binary outcomes, and 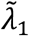 is the Lasso tuning parameter chosen by 3-fold cross-validation. Using the PRS as proxies for the effect estimates, we use the estimated ensembling weights 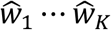 to obtain a final aggregated regression coefficient estimate:

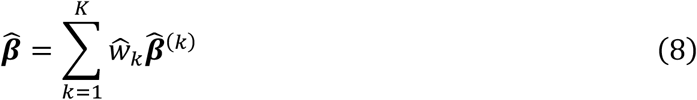

To get an insight into how this estimator improves over the traditional grid search-based estimator, using equation (6), the ensembled estimator 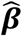 in equation (8) can be written as an iterative estimator

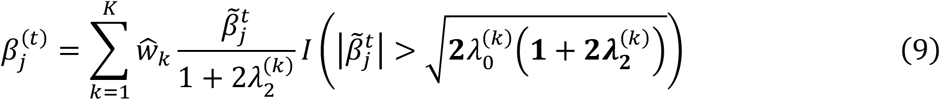

which combines the 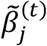 estimators obtained using different hard thresholds 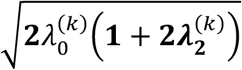 and shrinkages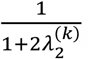. In fact, the traditional grid search corresponds to regressing ***Y***_*tun*_ on each 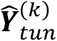 individually and identifying the single best 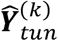 to predict ***Y***_*tun*_. This is equivalent to estimating the *w*_*k*_’s in the regression model of ***Y***_*tun*_ on 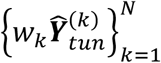 using a one-step forward selection procedure to identify the single best 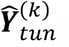, which will identify a model with inferior prediction performance compared to estimating *w*_*k*_’s by minimizing the problem in equation (7).

With the ensembled effect estimates, we construct a PRS in an out-of-sample validation dataset 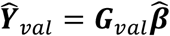, which remains separate from the cross-validation Lasso regression in the tuning dataset. Lastly, we assess the PRS’s prediction performance on the validation outcome ***Y***_*val*_, using R^2^ for continuous outcomes and AUC for binary outcomes.

### Implementation of PRS Methods

*C+T*^10,11^ is a method that first removes SNPs in high LD with an index SNP with the most significant GWAS p-value in a particular base-pair window. Then, several p-value thresholds are applied to the remaining SNPs to decide which parameter combination gives the best PRS. This was implemented in plink 1.9^54^ using LD thresholds *r*^2^ of 0.05 and 0.1, windows of size 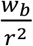 kb, where the window base *w*_*b*_ is either 25 or 50, and 23 p-value thresholds *p*^*T*^ ranging from 1x10^-11^ to 1. The use of 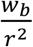 is motivated by an inverse relationship between LD and genetic distance between SNPs^91^. We choose the best clumping parameters and p-value threshold within the tuning dataset to construct the PRS.

*Stacked C+T*^28^ is an ensembling-based method that uses elastic net to aggregate PRS estimates from different clumping and thresholding parameterizations. Based on the online tutorial, we use 7 LD thresholds *r*^2^ between 0.01 and 0.95, 4 window bases *w*_M_ (50, 100, 200, and 500 kb), 50 p-value thresholds *p*^*T*^, and 3 weights *α* (1, 0.01, and 0.0001) for Lasso versus ridge.

*Lassosum2*^20^ is an extension of the previously developed Lassosum method^19^. The original Lassosum solves a Lasso regression using summary-level data from GWAS and LD, with an additional penalty applied to the LD matrix:

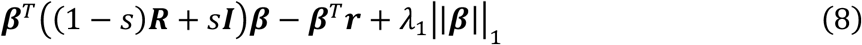

The tuning parameter *s* ∈ (0,1] is used to ensure convexity of the optimization, since the GWAS and LD are typically from different genotype data. The original Lassosum uses a block-diagonal LD approximation^41^ for ***R***, while Lassosum2 uses a banded LD approximation for each chromosome – the original results do not show much difference in prediction between the two LD approximations. For the grid search, we use 4 values of *s* (0.001, 0.01, 0.1, and 1) and 30 values of *λ*_1_.

*LDPred2*^14^ is a computational extension of the previously developed LDPred method^13^. This method is based on the Gaussian spike-and-slab prior for each SNP effect size *β*_*j*_:

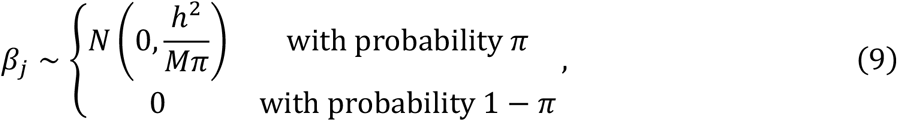

where *h*^2^ is the total heritability of a trait, *M* is the total number of SNPs, and *π* is the proportion of nonzero SNPs. Then, a Gibbs sampling MCMC algorithm is used to compute the posterior mean 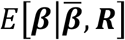, conditional on marginal effect sizes *ß* from GWAS and banded LD ***R***. LDPred2 introduces improvements in efficiency that allow for a larger grid search compared to LDPred. For this grid search, we use 0.7, 1, and 1.3 times the heritability *h*^2^ estimated from LD score regression^92^ – 21 values of *π* between 10^-5^ and 1, as well as “True” and “False” for the “sparse” setting that forces more posterior SNP effects to be 0.

Stacked C+T, Lassosum2, and LDPred2 were implemented using bigsnpr (version 1.8.1) in R, following online tutorials, with 10 cores for parallel computing^14,20,28,55^. For Lassosum2 and LDPred2, we used a 500 kb-width banded LD for each chromosome.

### Published GWAS Summary Statistics and Evaluation in UK Biobank

We further compare the PRS methods on 27 published summary statistics across 11 traits and 9 data sources. Tuning and validation were done using individual-level genotype and phenotype data from the UKBB^40^. We focus our analysis on individuals of European ancestry, based on self-reported race as “British”, “Irish”, “White”, or ‘Any other white background.” After excluding individuals who withdrew consent as of April 26, 2023, and have missing baseline measurements for age, sex, and the first ten PCs, and further restricting to unrelated samples so that no pair has relationship closer than fourth degree^93^, we have a total of 318,052 samples for analysis. 40,000 samples were randomly chosen for tuning and validation - with 20,000 samples in each split. We use the tuning sample as the LD reference for each of the PRS methods. Our analysis is restricted to about 1.5 million SNPs in HapMap3^49^ and Multi-Ethnic Genotyping Arrays (MEGA)^94^ chips that are shared between the GWAS and UKBB imputed genotype. Any remaining missing SNP values were assigned a dosage of 0 for consistency across methods. The 1.5 million SNPs were selected according to analysis of GLGC traits in the recently published CT-SLEB paper^29^. Note that quality control might result in varying number of SNPs across different datasets, but all PRS methods use the same GWAS summary statistics to ensure fair comparison.

The summary statistics for these traits were collected from a variety of publicly available sources (**Supplementary Tables 1-3**). Baseline covariates (age, sex, and the first ten PCs) and continuous trait measurements were obtained from the “initial assessment visit” period. Binary traits were defined using ICD10 codes, where the outcome is set as 1 if at least one code was observed and 0 otherwise (**Supplementary Table 4**). For tuning and validation of continuous traits, we compute R^2^ with respect to the residuals of the trait after regressing out the baseline covariates. For binary traits, we use the RISCA package (version 1.0.3) in R^60^ to compute adjusted AUC to account for baseline covariates in logistic regression.

**Main Figures**

## References

1. Chaoerjee, N., Shi, J. & García-Closas, M. Developing and evaluaIng polygenic risk predicIon models for straIfied disease prevenIon. Nat Rev Genet 17, 392–406 (2016).

2. Hung, R. J. et al. Assessing Lung Cancer Absolute Risk Trajectory Based on a Polygenic Risk Model. Cancer Res 81, 1607–1615 (2021).

3. Graham, S. E. et al. The power of geneIc diversity in genome-wide associaIon studies of lipids. Nature 600, 675–679 (2021).

4. Zhang, H. et al. Genome-wide associaIon study idenIfies 32 novel breast cancer suscepIbility loci from overall and subtype-specific analyses. Nat Genet 52, 572–581 (2020).

5. Wray, N. R. et al. Genome-wide associaIon analyses idenIfy 44 risk variants and refine the geneIc architecture of major depression. Nat Genet 50, 668–681 (2018).

6. Schaid, D. J., Sinnwell, J. P., Batzler, A. & McDonnell, S. K. Polygenic risk for prostate cancer: Decreasing relaIve risk with age but li0le impact on absolute risk. The American Journal of Human Gene:cs 109, 900–908 (2022).

7. Torkamani, A., Wineinger, N. E. & Topol, E. J. The personal and clinical uIlity of polygenic risk scores. Nature Reviews Genetics vol. 19 581–590 Preprint at 10.1038/s41576-018-0018-x (2018).

8. Lewis, C. M. & Vassos, E. Polygenic risk scores: From research tools to clinical instruments. Genome Medicine vol. 12 Preprint at 10.1186/s13073-020-00742-5 (2020).

9. Adeyemo, A. et al. Responsible use of polygenic risk scores in the clinic: potenIal benefits, risks and gaps. Nature Medicine vol. 27 1876–1884 Preprint at 10.1038/s41591-021-01549-6 (2021).

10. Wray, N. R., Goddard, M. E. & Visscher, P. M. PredicIon of individual geneIc risk to disease from genome-wide associaIon studies. Genome Res 17, 1520–1528 (2007).

11. Purcell, S. M. et al. Common polygenic variaIon contributes to risk of schizophrenia and bipolar disorder. Nature 460, 748–752 (2009).

12. Lo, A., Chernoff, H., Zheng, T. & Lo, S.-H. Why significant variables aren’t automaIcally good predictors. Proceedings of the National Academy of Sciences 112, 13892–13897 (2015).

13. Vilhjálmsson, B. J. et al. Modeling Linkage Disequilibrium Increases Accuracy of Polygenic Risk Scores. Am J Hum Genet 97, 576–592 (2015).

14. Privé, F., Arbel, J. & Vilhjálmsson, B. J. LDpred2: Be0er, faster, stronger. Bioinforma:cs 36, 5424–5431 (2020).

15. Lloyd-Jones, L. R. et al. Improved polygenic predicIon by Bayesian mulIple regression on summary staIsIcs. Nat Commun 10, (2019).

16. Ge, T., Chen, C. Y., Ni, Y., Feng, Y. C. A. & Smoller, J. W. Polygenic predicIon via Bayesian regression and conInuous shrinkage priors. Nat Commun 10, (2019).

17. Ni, G. et al. A Comparison of Ten Polygenic Score Methods for Psychiatric Disorders Applied Across MulIple Cohorts. Biol Psychiatry 90, 611–620 (2021).

18. Zhou, G. & Zhao, H. A fast and robust Bayesian nonparametric method for predicIon of complex traits using summary staIsIcs. PLoS Genet 17, e1009697 (2021).

19. Mak, T. S. H., Porsch, R. M., Choi, S. W., Zhou, X. & Sham, P. C. Polygenic scores via penalized regression on summary statistics. Genet Epidemiol 41, 469–480 (2017).

20. Privé, F., Arbel, J., Aschard, H. & Vilhjálmsson, B. J. Identifying and correcting for misspecifications in GWAS summary statistics and polygenic scores. Human Gene:cs and Genomics Advances 3, 100136 (2022).

21. Su, W., Bogdan, M. & Candès, E. False discoveries occur early on the Lasso path. The Annals of Sta:s:cs 45, (2017).

22. Slatkin, M. Linkage disequilibrium — understanding the evolutionary past and mapping the medical future. Nat Rev Genet 9, 477–485 (2008).

23. Liashchynskyi, P. & Liashchynskyi, P. Grid Search, Random Search, Genetic Algorithm: A Big Comparison for NAS. (2019).

24. Le, T. M. & Clarke, B. Model Averaging Is Asympto:cally BeGer Than Model Selec:on For Predic:on. Journal of Machine Learning Research vol. 23 (2022).

25. Jin, J. et al. ME-Bayes SL: Enhanced Bayesian Polygenic Risk Prediction Leveraging Information across Multiple Ancestry Groups. bioRxiv (2023) doi:10.1101/2023.04.12.536510.

26. Rauschenberger, A., Glaab, E. & Van De Wiel, M. A. Predictive and interpretable models via the stacked elastic net. Bioinformatics 37, 2012–2016 (2021).

27. van der Laan, M. J., Polley, E. C. & Hubbard, A. E. Super Learner. Stat Appl Genet Mol Biol 6, (2007).

28. Privé, F., Vilhjálmsson, B. J., Aschard, H. & Blum, M. G. B. Making the Most of Clumping and Thresholding for Polygenic Scores. Am J Hum Genet 105, 1213–1221 (2019).

29. Zhang, H. et al. A new method for multiancestry polygenic prediction improves performance across diverse populations. Nat Genet (2023) doi:10.1038/s41588-023-01501-z.

30. Zhang, J. et al. An Ensemble Penalized Regression Method for Multi-ancestry Polygenic Risk Prediction. bioRxiv (2023) doi:10.1101/2023.03.15.532652.

31. Hazimeh, H. & Mazumder, R. Fast best subset selection: Coordinate descent and local combinatorial optimization algorithms. Oper Res 68, 1517–1537 (2020).

32. Dedieu, A., Hazimeh, H. & Mazumder, R. Learning Sparse Classifiers: Con:nuous and Mixed Integer Op:miza:on Perspec:ves Dedieu, Hazimeh, and Mazumder. Journal of Machine Learning Research vol. 22 (2021).

33. Mazumder, R., Radchenko, P. & Dedieu, A. Subset Selection with Shrinkage: Sparse Linear Modeling When the SNR Is Low. Oper Res (2022) doi:10.1287/opre.2022.2276.

34. Khera, A. V. et al. Genome-wide polygenic scores for common diseases identify individuals with risk equivalent to monogenic mutations. Nat Genet 50, 1219–1224 (2018).

35. Wang, Z. et al. The Value of Rare Genetic Variation in the Prediction of Common Obesity in European Ancestry Populations. Front Endocrinol (Lausanne) 13, (2022).

36. Ellio, J. et al. Predictive Accuracy of a Polygenic Risk Score–Enhanced Prediction Model vs a Clinical Risk Score for Coronary Artery Disease. JAMA 323, 636 (2020).

37. Wong, C. K. et al. Polygenic risk scores for cardiovascular diseases and type 2 diabetes. PLoS One 17, e0278764 (2022).

38. Patel, A. P. et al. A multi-ancestry polygenic risk score improves risk prediction for coronary artery disease. Nat Med (2023) doi:10.1038/s41591-023-02429-x.

39. Kurki, M. I. et al. FinnGen provides genetic insights from a well-phenotyped isolated population. Nature 613, 508–518 (2023).

40. Sudlow, C. et al. UK Biobank: An Open Access Resource for Identifying the Causes of a Wide Range of Complex Diseases of Middle and Old Age. PLoS Med 12, e1001779 (2015).

41. Berisa, T. & Pickrell, J. K. Approximately independent linkage disequilibrium blocks in human populations. Bioinformatics 32, 283–285 (2016).

42. Polson, N. G. & Sun, L. Bayesian L0-regularized least squares. Appl Stoch Models Bus Ind 35, 717–731 (2019).

43. Park, J.-H. et al. Distribution of allele frequencies and effect sizes and their interrelationships for common genetic susceptibility variants. Proceedings of the National Academy of Sciences 108, 18026–18031 (2011).

44. Loh, P.-R. et al. Efficient Bayesian mixed-model analysis increases association power in large cohorts. Nat Genet 47, 284–290 (2015).

45. Henderson, C. R. Best Linear Unbiased Estimation and Prediction under a Selection Model. Biometrics 31, 423 (1975).

46. Meuwissen, T. H. E., Hayes, B. J. & Goddard, M. E. Prediction of Total Genetic Value Using Genome-Wide Dense Marker Maps. Genetics 157, 1819–1829 (2001).

47. Friedman, J., Hastie, T. & Tibshirani, R. Regularization Paths for Generalized Linear Models via Coordinate Descent. JSS Journal of Sta:s:cal SoSware vol. 33 https://www.jstatsos.org/ (2010).

48. Su, Z., Marchini, J. & Donnelly, P. HAPGEN2: Simulation of multiple disease SNPs. Bioinformatics 27, 2304–2305 (2011).

49. International HapMap 3 Consortium. Integrating common and rare genetic variation in diverse human populations. Nature 467, 52–58 (2010).

50. Yang, J., Lee, S. H., Goddard, M. E. & Visscher, P. M. GCTA: A Tool for Genome-wide Complex Trait Analysis. The American Journal of Human Genetics 88, 76–82 (2011).

51. Daetwyler, H. D., Villanueva, B. & Woolliams, J. A. Accuracy of predicting the genetic risk of disease using a genome-wide approach. PLoS One 3, (2008).

52. Wray, N. R. et al. Pitalls of predicting complex traits from SNPs. Nature Reviews Genetics vol. 14 507–515 Preprint at 10.1038/nrg3457 (2013).

53. Ding, Y. et al. Large uncertainty in individual polygenic risk score estimation impacts PRS-based risk stratification. Nat Genet 54, 30–39 (2022).

54. Chang, C. C. et al. Second-generation PLINK: rising to the challenge of larger and richer datasets. Gigascience 4, 7 (2015).

55. Privé, F., Aschard, H., Ziyatdinov, A. & Blum, M. G. B. Efficient analysis of large-scale genome-wide data with two R packages: bigstatsr and bigsnpr. Bioinformatics 34, 2781–2787 (2018).

56. Demenais, F. et al. Multiancestry association study identifies new asthma risk loci that colocalize with immune-cell enhancer marks. Nat Genet 50, 42–53 (2018).

57. Nikpay, M. et al. A comprehensive 1000 Genomes-based genome-wide association metaanalysis of coronary artery disease. Nat Genet 47, 1121–1130 (2015).

58. Mahajan, A. et al. Fine-mapping type 2 diabetes loci to single-variant resolution using high-density imputation and islet-specific epigenome maps. Nat Genet 50, 1505–1513 (2018).

59. Schumacher, F. R. et al. Association analyses of more than 140,000 men identify 63 new prostate cancer susceptibility loci. Nat Genet 50, 928–936 (2018).

60. Foucher, Y., Le Borgne, F., Chaoon, A. & Sabathe, C. RISCA: Causal Inference and Prediction in Cohort-Based Analyses. Preprint at https://CRAN.R-project.org/package=RISCA (2022).

61. Sun, B. B. et al. Genetic associations of protein-coding variants in human disease. Nature 603, 95–102 (2022).

62. Auton, A. et al. A global reference for human genetic variation. Nature 526, 68–74 (2015).

63. Schoeler, T. et al. Participation bias in the UK Biobank distorts genetic associations and downstream analyses. Nat Hum Behav (2023) doi:10.1038/s41562-023-01579-9.

64. Lewis, A. C. F. & Green, R. C. Polygenic risk scores in the clinic: new perspectives needed on familiar ethical issues. Genome Med 13, (2021).

65. Hao, L. et al. Development of a clinical polygenic risk score assay and reporting workflow. Nat Med 28, 1006–1013 (2022).

66. Krainc, T. & Fuentes, A. Genetic ancestry in precision medicine is reshaping the race debate. Proceedings of the National Academy of Sciences 119, (2022).

67. Ding, Y. et al. Polygenic scoring accuracy varies across the genetic ancestry continuum. Nature (2023) doi:10.1038/s41586-023-06079-4.

68. Pärna, K. et al. A Principal Component Informed Approach to Address Polygenic Risk Score Transferability Across European Cohorts. Front Genet 13, (2022).

69. Lennon, N. J. et al. Selection, optimization, and validation of ten chronic disease polygenic risk scores for clinical implementation in diverse populations. medRxiv (2023) doi:10.1101/2023.05.25.23290535.

70. Cai, M. et al. A unified framework for cross-population trait prediction by leveraging the genetic correlation of polygenic traits. Am J Hum Genet 108, 632–655 (2021).

71. Albiñana, C. et al. Leveraging both individual-level genetic data and GWAS summary statistics increases polygenic prediction. Am J Hum Genet 108, 1001–1011 (2021).

72. Weissbrod, O. et al. Leveraging fine-mapping and multipopulation training data to improve cross-population polygenic risk scores. Nat Genet 54, 450–458 (2022).

73. Ruan, Y. et al. Improving polygenic prediction in ancestrally diverse populations. Nat Genet 54, 573–580 (2022).

74. Zhao, Z., Fritsche, L. G., Smith, J. A., Mukherjee, B. & Lee, S. The Construction of Multi-ethnic Polygenic Risk Score using Transfer Learning. doi:10.1101/2022.03.08.22272114.

75. Sun, Q. et al. Improving polygenic risk prediction in admixed populations by explicitly modeling ancestral-specific effects via GAUDI. doi:10.1101/2022.10.06.511219.

76. Machado Reyes, D., Bose, A., Karavani, E. & Parida, L. FairPRS: adjus:ng for admixed popula:ons in polygenic risk scores using invariant risk minimiza:on. Error! Hyperlink reference not valid.(2022).

77. Kachuri, L. et al. Principles and methods for transferring polygenic risk scores across global populations. Nat Rev Genet (2023) doi:10.1038/s41576-023-00637-2.

78. Kim, J. & Park, H. Fast Active-set-type Algorithms for L1-regularized Linear Regression. Interna:onal Conference on Ar:ficial Intelligence and Sta:s:cs (2010).

79. Nutini, J., Laradji, I. & Schmidt, M. Let’s Make Block Coordinate Descent Converge Faster: Faster Greedy Rules, Message-Passing, Active-Set Complexity, and Superlinear Convergence. Journal of Machine Learning Research 23, 1–74 (2022).

80. Polley, E., LeDell, E., Kennedy, C. & van der Laan, M. SuperLearner: Super Learner Prediction. Preprint at https://CRAN.R-project.org/package=SuperLearner (2021).

81. Zhao, Z. et al. PUMAS: fine-tuning polygenic risk scores with GWAS summary statistics. Genome Biol 22, (2021).

82. Zhao, Z. et al. Optimizing and benchmarking polygenic risk scores with GWAS summary statistics. bioRxiv (2022) doi:10.1101/2022.10.26.513833.

83. Miao, J. et al. Quantifying portable genetic effects and improving cross-ancestry genetic prediction with GWAS summary statistics. Nat Commun 14, 832 (2023).

84. Coombes, B. J., Ploner, A., Bergen, S. E. & Biernacka, J. M. A principal component approach to improve association testing with polygenic risk scores. Genet Epidemiol 44, 676–686 (2020).

85. Jiang, W., Chen, L., Girgenti, M., Zhao, H. & Girgenti, M. J. Tuning Parameters for Polygenic Risk Score Methods Using GWAS Summary Statistics from Training Data Tuning Parameters for Polygenic Risk Score. (2023) doi:10.21203/rs.3.rs-2939390/v1.

86. Zou, H. & Hastie, T. Regulariza:on and variable selec:on via the elas:c net. J. R. Statist. Soc. B vol. 67 (2005).

87. Tibshirani, R. Regression Shrinkage and Selection via the Lasso. Source: Journal of the Royal Sta:s:cal Society. Series B (Methodological) vol. 58 (1996).

88. Bertsimas, D., King, A. & Mazumder, R. Best subset selection via a modern optimization lens. Ann Stat 44, 813–852 (2016).

89. Fan, J., Liao, Y. & Mincheva, M. High-dimensional covariance matrix estimation in approximate factor models. The Annals of Sta:s:cs 39, (2011).

90. Gao, J., Han, X., Pan, G. & Yang, Y. High dimensional correlation matrices: the central limit theorem and its applications. J R Stat Soc Series B Stat Methodol 79, 677–693 (2017).

91. Pritchard, J. K. & Przeworski, M. Linkage Disequilibrium in Humans: Models and Data. The American Journal of Human Genetics 69, 1–14 (2001).

92. Bulik-Sullivan, B. et al. LD score regression distinguishes confounding from polygenicity in genome-wide association studies. Nat Genet 47, 291–295 (2015).

93. Dey, R. et al. Efficient and accurate frailty model approach for genome-wide survival association analysis in large-scale biobanks. Nat Commun 13, 5437 (2022).

94. Bien, S. A. et al. Strategies for Enriching Variant Coverage in Candidate Disease Loci on a Multiethnic Genotyping Array. PLoS One 11, e0167758 (2016).

