## Supplementary Figures and Method Notes for "Ensembled best subset selection using summary statistics for polygenic risk prediction"

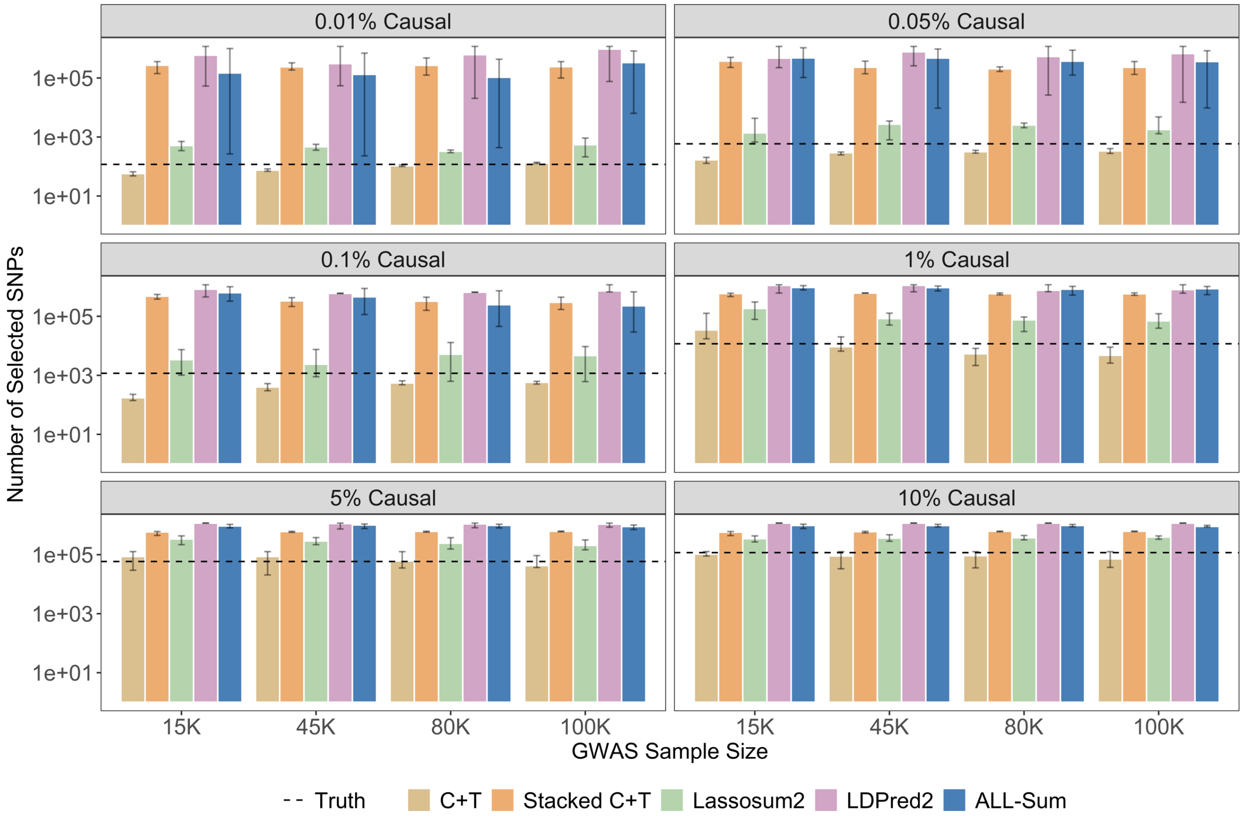


**Supplementary Figure 1 Comparison of PRS sparsity in simulation studies.** We compare the number of selected SNPs (Y-axis, log scale) in the final PRS across the 6 causal SNP proportions (panels) and 4 GWAS sample sizes (X-axis). Methods like C+T and Lassosum2 are based on sparse modeling and grid search, and intuitively select a relatively small number of SNPs on a similar scale as the true number of causal SNPs. Bayesian methods like LDPred2 use posterior averaging to get estimates, and thus tend to select a very large number of SNPs (nearly all 1.2 million) despite the incorporation of sparsity in their priors. Stacked C+T and ALL-Sum perform model averaging over many sparse estimates – C+T and $L_0L_2$ regression, respectively – so the final estimates also select a fairly large number of SNPs, smaller than or similar to LDPred2.

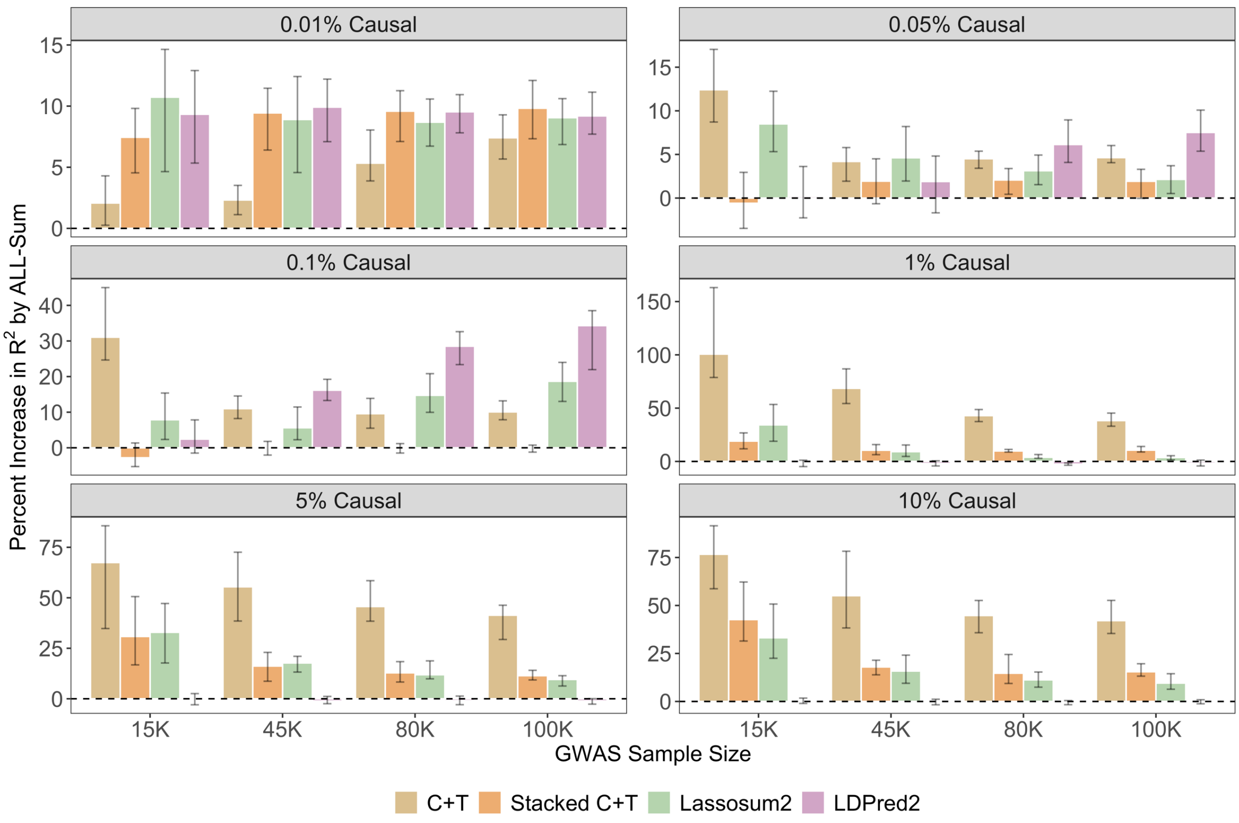


**Supplementary Figure 2 Improvement in prediction accuracy by ALL-Sum over other PRS methods in simulation studies.** We compare the percentage change in $R^2$ (Y-axis) of ALL-Sum from other PRS methods across the 6 causal SNP proportions (panels) and 4 GWAS sample sizes (X-axis). Bar plots show the average percentage change over 10 simulation replicates, and error bars denote the range. For nearly every simulation setting, ALL-Sum shows a large percentage increase in $R^2$ over all four alternative PRS methods. In a few scenarios, ALL-Sum performs about comparable to either Stacked C+T or LDPred2.

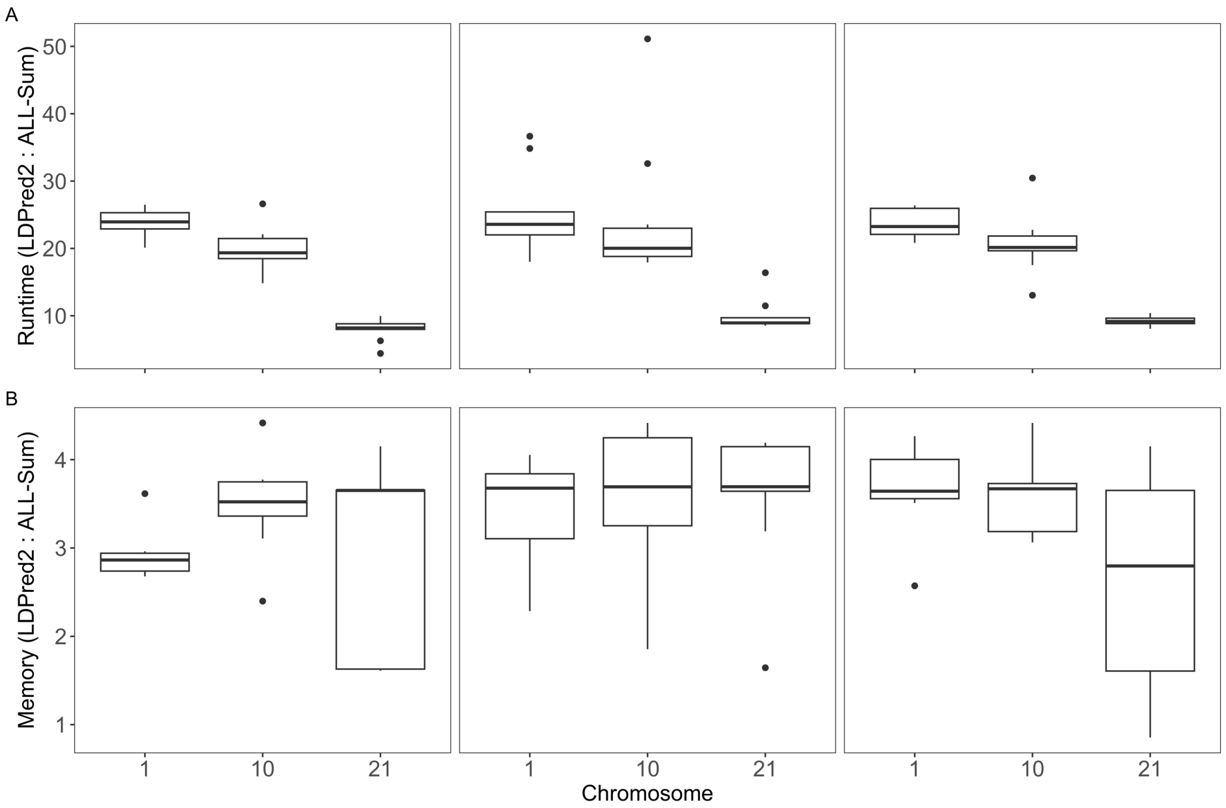


**Supplementary Figure 3 Relative runtime and memory usage comparison between ALL-Sum and LDPred2**. We measure the ratio in runtime (Y-axis A) and memory usage (Y-axis B) of LDPred2 relative to ALL-Sum using simulated data on chromosomes 1, 10, and 21 (X-axis), with causal SNP proportions 0.1%, 1%, and 10% (panels) and GWAS sample size of 100,000.

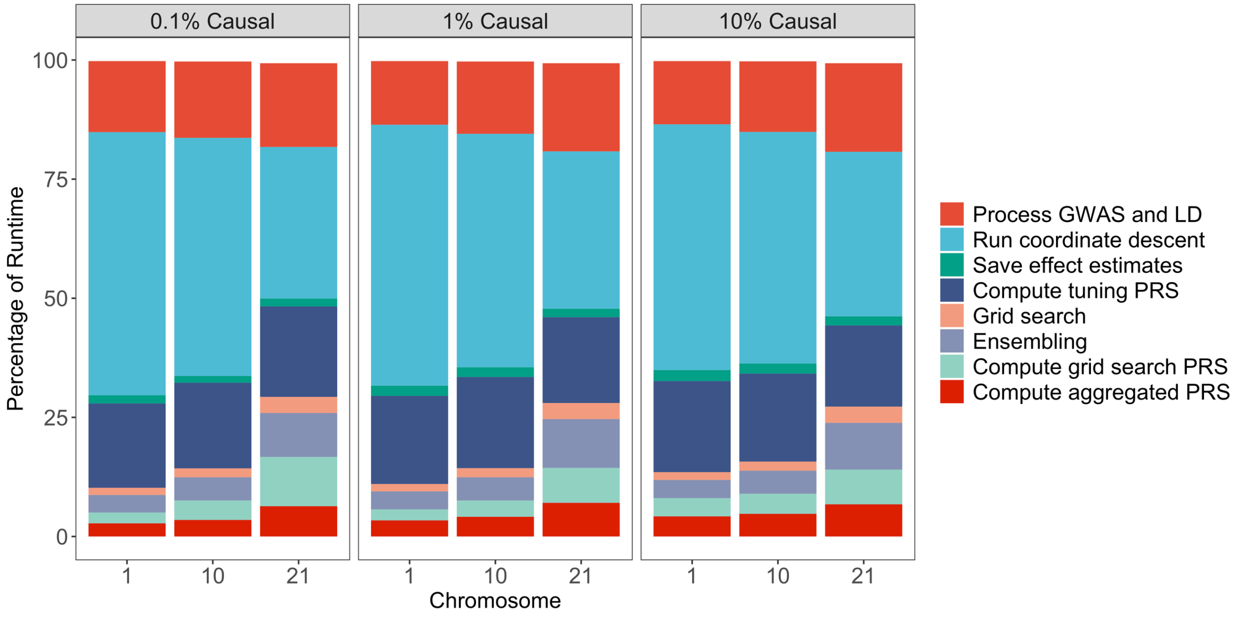


**Supplementary Figure 4 Runtime distribution for ALL-Sum**. We compare the percentage of runtime for each stage of the ALL-Sum procedure (Y-axis) for 3 causal SNP proportions (panels) and 3 chromosomes (X-axis). We can see that runtime is consistent across causal SNP proportions, but the distribution shifts as the dimension increases from chromosome 21 (16,865 SNPs) to 1 (97,230 SNPs). Most runtime is taken by the coordinate descent step (light blue), although for smaller-dimension problems, a slightly larger proportion of time is taken up by computing PRS at the tuning and validation steps (dark blue, light green, dark red).

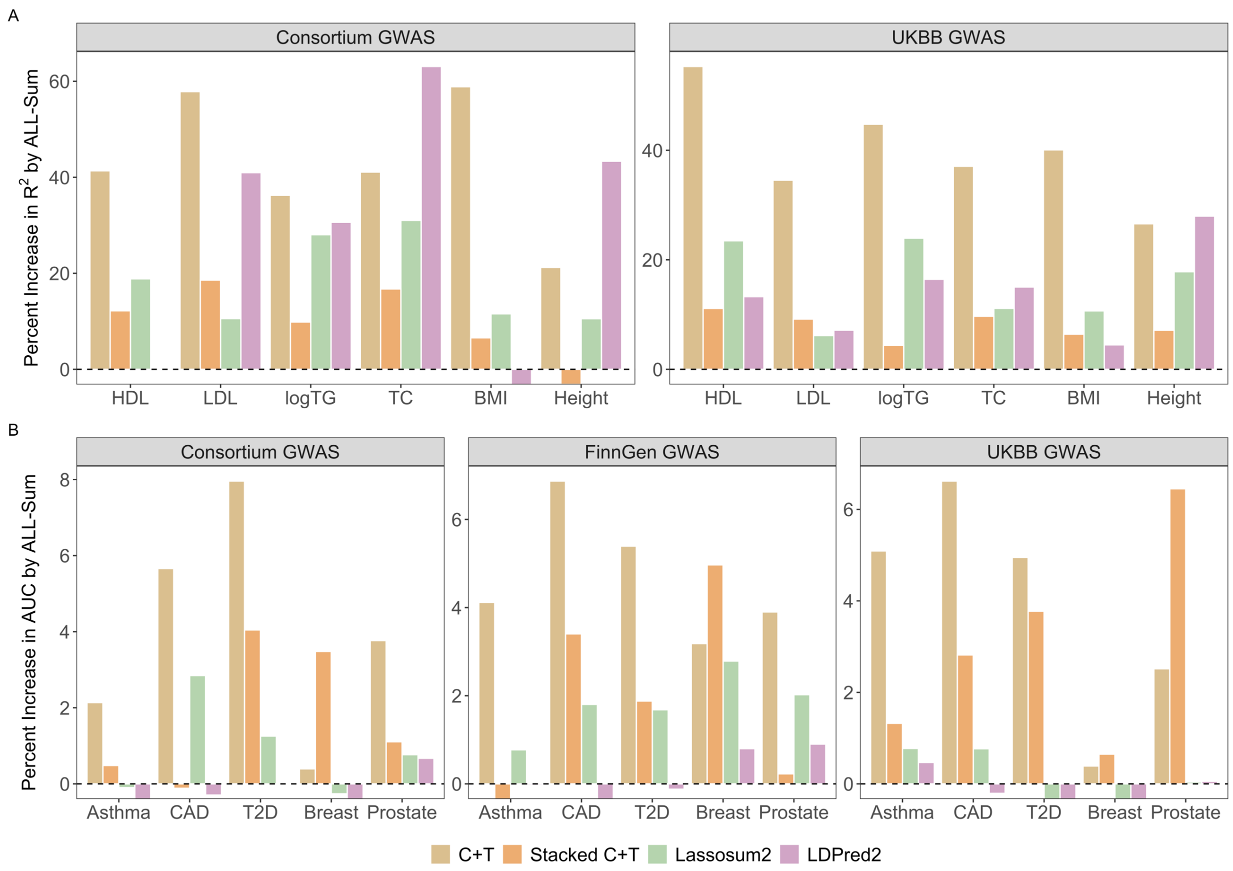


**Supplementary Figure 5 Improvement in prediction accuracy by ALL-Sum over other PRS methods in real continuous and binary traits** We compare the percentage change in $R^2$ for continuous traits (Y-axis A) or AUC for binary traits (Y-axis B) of ALL-Sum from other PRS methods for either consortium-, FinnGen-, or UKBB-based GWAS summary statistics. For nearly every trait and data source, ALL-Sum shows a large percentage increase in $R^2$ over all four alternative PRS methods.

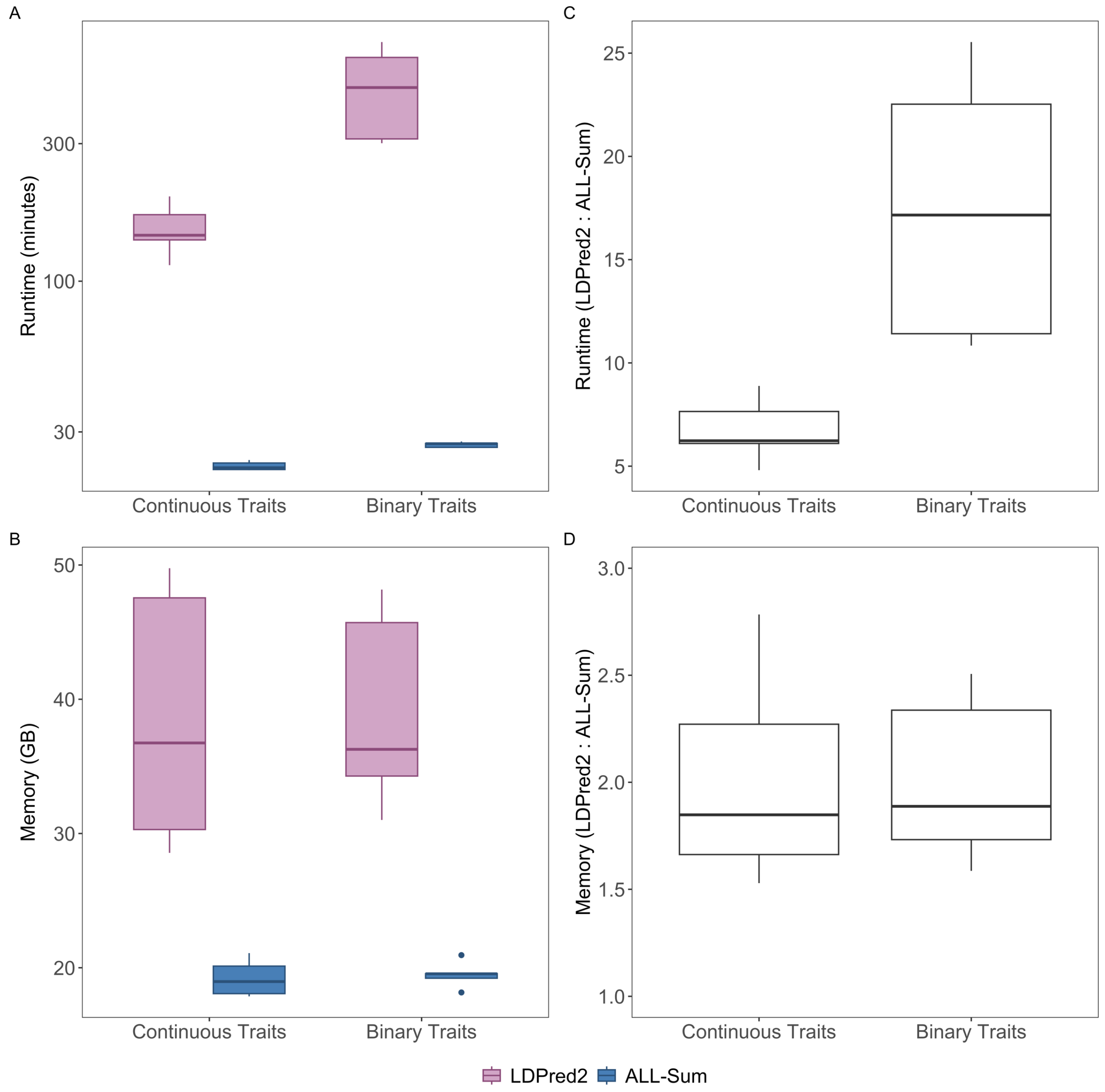


**Supplementary Figure 6 Runtime and memory usage comparison ALL-Sum and LDPred2 using UKBB-based GWAS summary statistics**. ALL-Sum and LDPred2 were run on full-genome summary statistics (about 1.5 million SNPs and 280,000 samples). We evaluate the runtime (Y-axis A) and memory usage (Y-axis B) of the full analysis procedure with no parallelization, as well as ratios for the relative increase in runtime (Y-axis C) and memory usage (Y-axis D) by LDPred2.

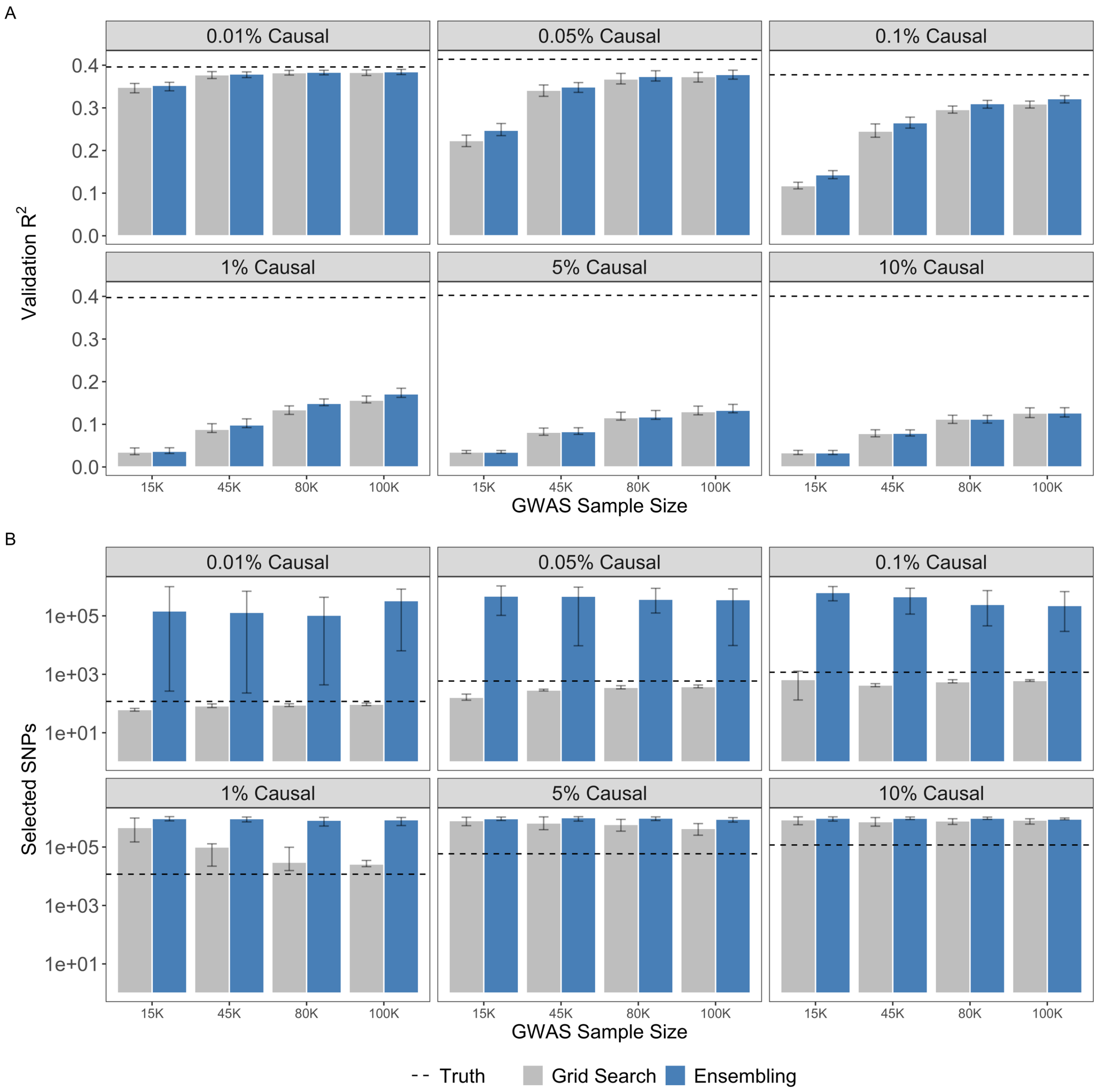


**Supplementary Figure 7 Comparison of grid search and ensembling in simulation studies.** We evaluate the prediction accuracy (Y-axis A) and number of selected SNPs (Y-axis B) of using tuning parameter grid search or ensembling to choose the best PRS after $L_0L_2$ penalized regression using the L0Learn-Sum optimization.

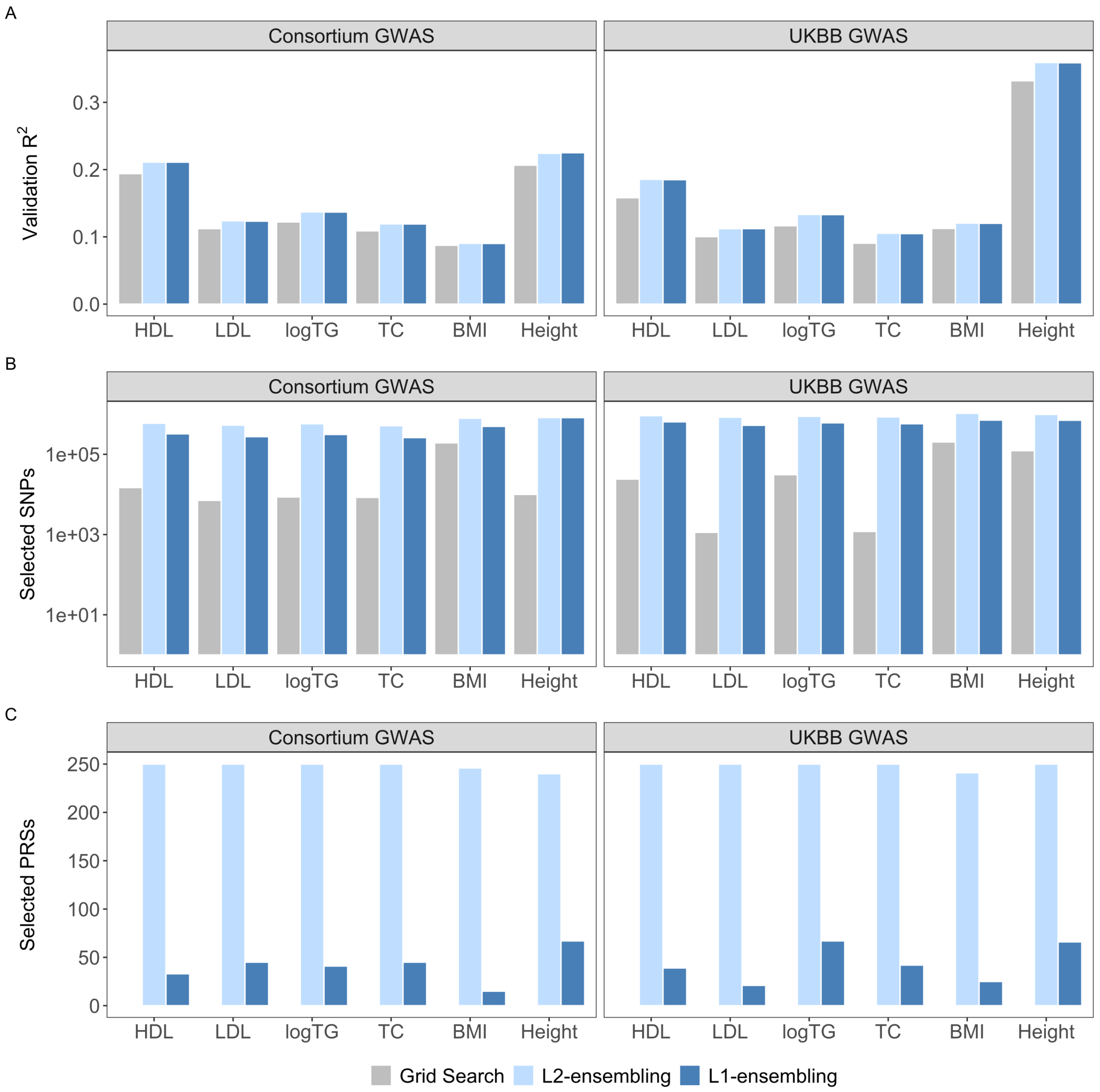


**Supplementary Figure 8 Comparison of ensembling strategies in continuous traits**. We compare the prediction performance (Y-axis A), number of selected SNPs (Y-axis B), and number of selected PRS (Y-axis C) of grid search of tuning parameters as well as ridge (L2) and lasso (L1) to ensemble the regression coefficients estimated using different grids, across 6 continuous traits (X-axis) and either consortium-based or UKBB-based GWAS summary statistics (panels).

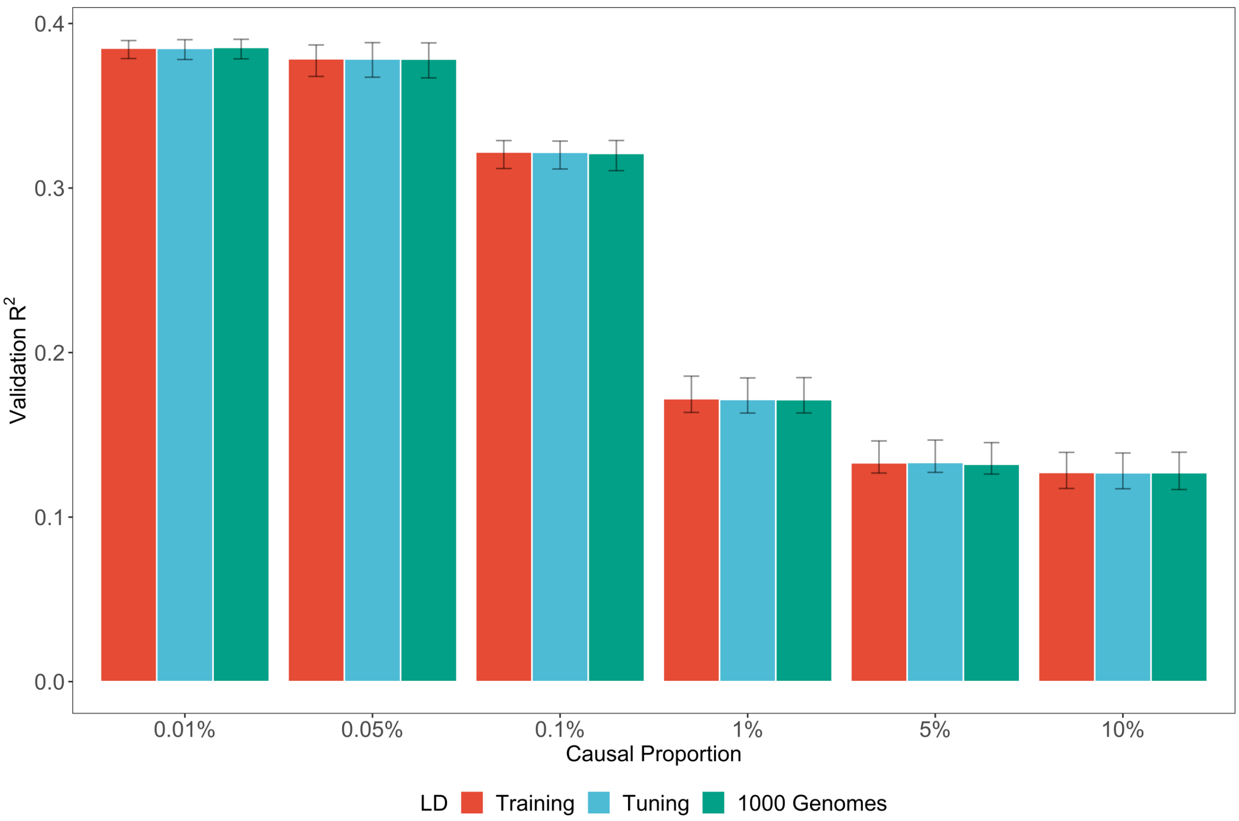


**Supplementary Figure 9 Comparison of prediction performance using different LD references in simulation studies**. We compare the prediction accuracy (Y-axis) of ALL-Sum using three different LD references: 100,000 training samples used to run the original GWAS, 10,000 tuning samples as in the main analysis, and 503 European samples from the 1000 Genomes Projects. We evaluate performance using the 6 different causal SNP proportions (X-axis) restricted to the 100,000 GWAS sample size case.

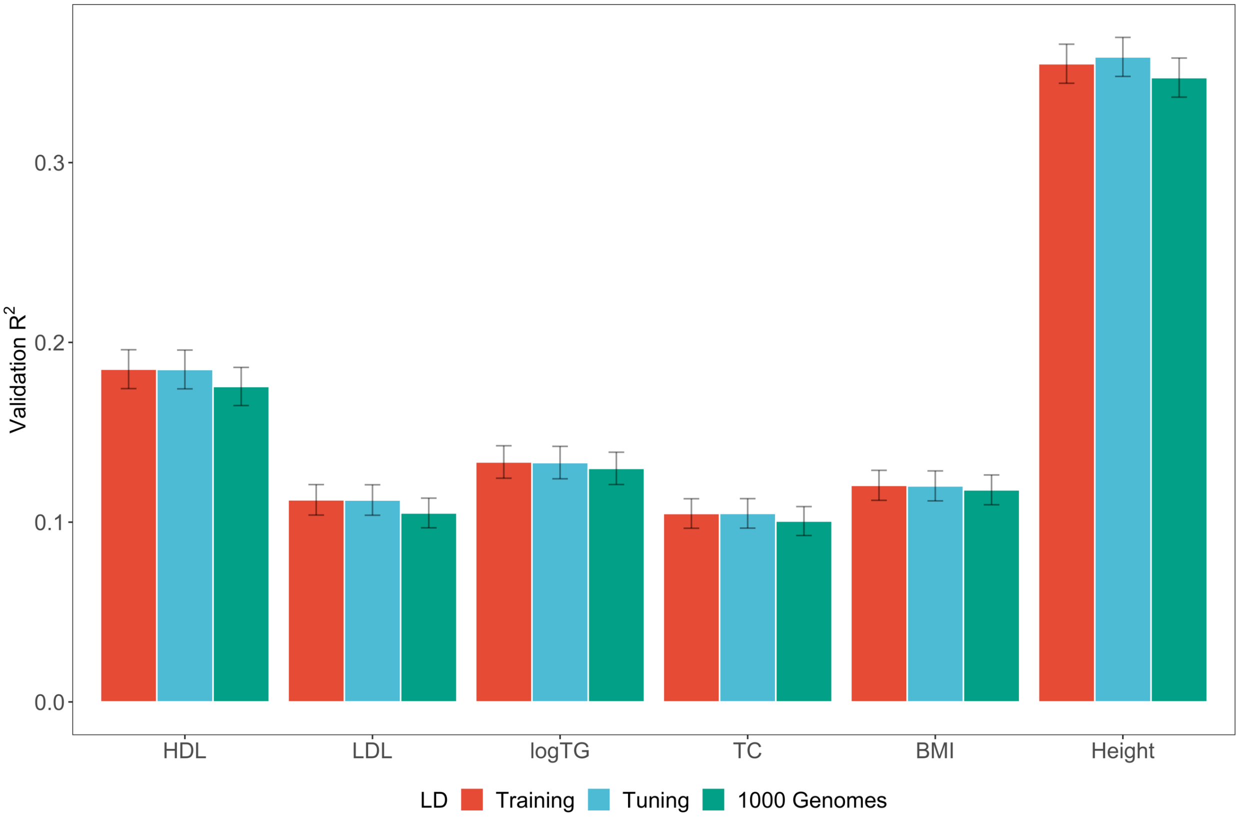


**Supplementary Figure 10 Comparison of prediction performance using different LD references for UKBB-based continuous trait analysis**. We compare the prediction accuracy (Y-axis) of ALL-Sum using three different LD references: 278,052 training samples used to run the original GWAS, 20,000 tuning samples as in the main analysis, and 503 European samples from the 1000 Genomes Project. We evaluate performance on the 4 lipid traits (X-axis) using UKBB-based GWAS.

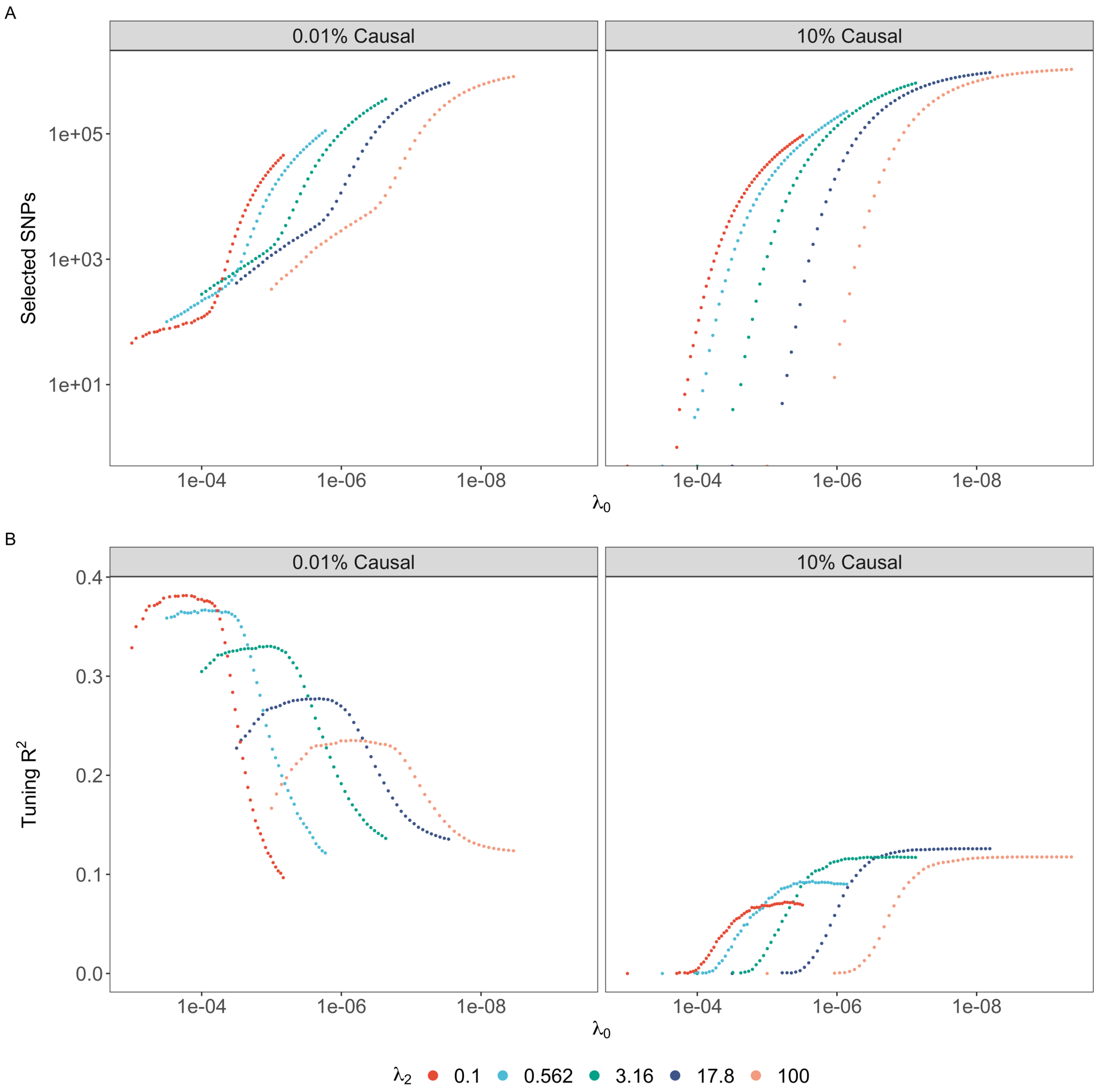


**Supplementary Figure 11 Demonstration of adaptive regularization path along $\lambda_0$ with varying $\lambda_2$ in simulated data**. Using an adaptive sequence of $\lambda_0$ (X-axis) for various chosen values of $\lambda_2$ (coloring) ensures that the number of selected SNPs (Y-axis A) is different for each solution in the tuning parameter grid. Furthermore, the best parameter combination based on $R^2$ in a tuning sample (Y-axis B) can be identified within the interior of the grid for extremely low and high causal SNP proportions (panels), verifying the flexibility of the grid specifications.

### Supplementary Method Notes

#### *Partial sorting of summary statistics*

The coordinate descent algorithm is a simple and fast algorithm on its own, but to further boost efficiency, we do “partial sorting” before running the optimization. In this step, we take an initial scan through all SNPs and order them from largest to smallest values of $|r_j|$. With the block-diagonal approximation, this sorting is thus done in blocks. This “greedy” approach prioritizes important SNPs in coordinate descent, which in turn helps the algorithm obtain better estimates and reach convergence faster[31,79].

#### *Adaptive regularization path along $\lambda_0$*

Most PRS methods require a grid search across several different combinations of tuning parameters to approximately achieve the best out-of-sample prediction. For $L_0L_2$ regression, we can choose a log-scale vector of $\lambda_2$ values to get broad coverage of varying levels of shrinkage. Then, most simply, for each value of $\lambda_2$ we can also set a fixed log-scale range of $\lambda_0$. However, since we are solving the penalized version of the best subset selection problem, we cannot directly control the number of nonzero SNPs using $\lambda_0$ like we could with the original constrained optimization problem[31,88]. As a result, different values of $\lambda_0$ can result in virtually identical solutions. Therefore, it is desirable to use a sequence of $\lambda_0$ values that yields different solutions for each $\lambda_0$ to make the most out of the grid (**Supplementary Figure 11**). This “adaptive regularization path” is again modified from the original L0Learn algorithm to accommodate for summary-level inputs.

For a fixed value of $\lambda_2$ and an initial $\lambda_0^{(0)}$, after getting the $k$-th solution $\widehat{\boldsymbol{\beta}}^{(k)}$, we can choose the subsequent value of $\lambda_0 = \lambda_0^{(k+1)}$ that guarantees at least one new nonzero entry in $\widehat{\boldsymbol{\beta}}^{(k+1)}$. First, we compute an upper bound derived from a reverse-engineering of the coordinate descent thresholding:

$$M^{(k)} = \frac{1}{2(1+2\lambda_2)} \max_{j:\widehat{\beta}_j^{(k)}=0} \left(\left|r_j - R_{j,\cdot}^T \hat{\beta}^{(k)}\right| - \lambda_1\right)^2 \qquad (S1)$$

By this construction, we know that $M^{(k)} < \lambda_0^{(k)}$. To ensure that we fully explore the space of candidate solutions, we can scale down this upper bound by a factor $\alpha$ between 0 and 1:

$$\lambda_0^{(k+1)} = \alpha M^{(k)} \qquad (S2)$$

Choosing an appropriate value of $\alpha$ poses a trade-off between thoroughly exploring the solution space and fine-tuning for the “best” solution. In our implementation of ALL-Sum, we let $\alpha$ vary inversely with $\lambda_2$. When $\lambda_2$ is large we expect effect sizes to be denser and smaller, and thus using a smaller $\alpha$ can allow a larger number of new SNPs to enter the model with the next $\lambda_0$ value. Conversely, when $\lambda_2$ is small, we expect sparser effect sizes, in which case a larger value of $\alpha$ limits the number of new selected SNPs to focus on sparse solutions. In the context of ensembling, this ensures that we have a wide range of candidate PRSs from which we can build a flexible final prediction model.
